# Peripheral vascular strand development in nodules is controlled by a bHLH/HLH heterodimer

**DOI:** 10.1101/2023.07.01.547324

**Authors:** Deevita Srivastava, Asim Ghosh, Michael Udvardi, Aashish Ranjan, Senjuti Sinharoy

## Abstract

Only the Leguminosae family can develop root nodules with peripheral vasculature, an adaptation that grants them an advantage in optimizing nitrogen fixation efficiency. *Medicago truncatula* develops indeterminate nodules that possess peripheral vascular-strands encircling the central infection zone. How vascular-strands shifted from the nodule central part to the periphery remains unresolved. Here we show, MtbHLH1 (renamed as Nodule Vascular bundle Development 1) is required for the proper organization of vascular strands. In *nvd1* nodules, vascular strands pass through the infection zone. *NVD2*, an HLH transcription factor that lacks a DNA-binding domain, is activated by NVD1. Mutant *nvd2* nodules display a similar partially central vasculature. *NVD2* is expressed along the nodule vascular bundle and NVD2:GFP fusion protein localizes to the nodule vascular endodermis. The formation of the peripheral vasculature is dependent on the proper stoichiometry of NVD1 and NVD2 heterodimers, as NVD2 controls NVD1-mediated transcriptional activation by sequestering NVD1. Transcription of *NVD1* is activated by auxin and Auxin Responsive transcription Factor (MtARF5). Transcriptome sequencing of *nvd1* and *nvd2* nodules and visualization of *in situ* auxin and cytokinin signal outputs indicated aberrant auxin/cytokinin balance in these nodules. Our findings showed that the NVD1-NVD2 heterodimer plays a key role in the formation of an orderly peripheral vascular bundle around Medicago nodules.

## Introduction

Analysis of molecular phylogeny has indicated that nitrogen-fixing root nodule symbiosis (RNS) is restricted to a monophyletic group of angiosperm order, the Fabids, consistent with a single origin of RNS, in a common ancestor of the Leguminosae and non-legume root nodule-forming families (Werner et al., 2014; Griesmann et al., 2018). Nitrogen-fixing non-legumes interact with actinorhizal bacteria to form nodules with a single vascular bundle (VB) in the center – like that of lateral roots. In contrast, legumes interact with rhizobia to develop nodules with multiple peripheral vasculatures (Pawlowski and Sprent, 2008; Raul et al., 2019). The evolution of peripheral vasculature in nodules is believed to have increased the nitrogen fixation efficiency of legumes compared to non-legumes by enabling the formation of a protective oxygen barrier around nitrogen-fixing cells of the nodule core, facilitating nutrient exchange, and/or serving as a major ATP-generating source (Downie, 2014; Venado et al., 2022).

In *M. truncatula*, Nod(ulation) factors (NF) signaling activates cell divisions at the pericycle and two innermost cortical cell layers within 24 hours post-inoculation (hpi). Further, endodermis and outer cortex-derived cells started dividing around 42-48 hpi. The dividing inner cortical cells become infected; however, the pericycle and endodermis-derived cells generate the nodule VB and remain uninfected (Ting Xiao et al., 2014). Medicago forms a cylindrical-shaped indeterminant nodule, with a) a persistent nodule meristem (NM) in the apex, followed by four distinct consecutive zones; b) an invasion zone (ZII); c) Interzone (ZII-ZIII); d) nitrogen-fixing zone; and e) senescence zone - only present in old nodules (Vasse et al., 1990). The NM is composed of different cell types in the central and peripheral regions. The central part of NM is marked by high cytokinin (CK) and is called nodule central meristem (NCM). NCM adds the central infection zone to a nodule (Ting Xiao et al., 2014; Triozzi et al., 2022). Nodule vascular meristems (NVMs), are marked by high auxin levels, are located on the edges of the NM, and are responsible for adding VB to a developing nodule (Franssen et al., 2015). *M. truncatula* NOD FACTOR HYDROLASE 1 (NFH1) hydrolyzes NF and thereby controls NF level. Studies have shown that the overexpression of *MtNFH1* or nodules of *Mtnfh1* and *Mtcre1* (*cytokinin receptor gene*) mutant plants developed multi-lobed nodules as opposed to the cylindrical structure which contains multiple NMs (Plet et al., 2011).

CK is necessary and sufficient for nodule organogenesis. The exogenous application of CK stimulates the formation of pseudo-nodules with peripheral vasculatures only in naturally nodule-forming legumes, suggesting that CK responsiveness of legume roots plays a pivotal role in the evolution of nodules with multiple peripheral vasculatures (Gauthier-Coles et al., 2018; Liu and Bisseling, 2020). Gene encoding the first and rate-limiting enzyme of CK biosynthesis pathway *adenylate isopentenyl transferase* (*IPT1/4*) and CK-activating enzyme *LONELY GUY* (*LOG*) express significantly during the *M. truncatula* nodule organogenesis (Gonzalez-Rizzo et al., 2006; Mortier et al., 2014; Azarakhsh et al., 2015; Van Zeijl et al., 2015). During nodule development, the CK signal is detected through MtCRE1. Two type-A response regulators, *MtRR11* and *MtRR4*, are activated early in the process (Gonzalez-Rizzo et al., 2006; Plet et al., 2011). *MtRR4* is activated by the ETHYLENE RESPONSE FACTOR REQUIRED FOR NODULE DIFFERENTIATION (EFD1) (Jardinaud et al., 2022). The type B response regulator *MtRR1* is actively transcribed in the susceptible zone of the root pericycle and is presumably the main candidate for CK signaling-mediated activation of the central regulator of nodule development, *NODULE INCEPTION* (*NIN*) (Liu et al., 2019). NIN is a critical point of divergence in the lateral root developmental program that promotes nodule development by directly activating three important transcription factors (Soyano et al., 2013; Baudin et al., 2015; Schiessl et al., 2019; Soyano et al., 2019). The NIN signaling pathway also activates the expression of auxin biosynthesis genes, specifically *MtYUC2* and 8 (Schiessl et al., 2019). Furthermore, the auxin influx carrier gene *MtLAX2* from the AUX/LAX family plays a vital role in nodule initiation by increasing auxin concentration (Roy et al., 2017).

Laser-capture microdissection (LCM)-based transcriptomes of different nodule zones have identified activation of auxin- and CK-induced genes in the nodule apex (Roux et al., 2014). The expression of important regulators of the root apical meristem (RAM) showed that certain genes specific to the RAM were predominantly expressed in the nodule vascular meristem (NVM), while others were expressed in the nodule cortical meristem (NCM). Indicating a reminiscent root developmental program is present in the NVM (Franssen et al., 2015). GRAS family transcription factors *SCARECROW* (*SCR*) and *MtEFD1* express at the NVM (Roux et al., 2014). The nodule identity is regulated by two paralogs: NODULE ROOT1 (MtNOOT1) and MtNOOT2, which suppress the root developmental program at the nodule apex and express in NVM and NCM respectively (Couzigou et al., 2012; Magne et al., 2018). *SHORT-ROOT* (*SHR*) and *SCARECROW* (*SCR*) are required for the establishment of the root endodermal cell layer (Shaar-Moshe and Brady, 2022). CK treatment and inoculation with rhizobia promote the accumulation of MtSHR1 and induce the expression of *MtSCR*. MtSHR1 promotes root cortical cell division downstream of NF-signaling. MtSHR1/2-mediated cortical cell division during nodule development is dependent upon SCR, and three nodule-specific transcription factors NODULATION SIGNALING PATHWAY1 (NSP1), NSP2, NIN (Dong et al., 2021). In Arabidopsis, *SCR* expression is also regulated by another transcription factor JACKDAW (Welch et al., 2007)*. MtSHR1/2* and *MtJACKDAW (JKD)* are also expressed in the nodule apex (Schlereth et al., 2010; Roux et al., 2014).

Our knowledge is limited regarding the evolution of peripheral VB development in legumes. *M. truncatula lumpy infections (lin-1/4),* and *vapyrin-2* nodules are small, devoid of infection, and have central VB. Similarly, an *Sinorhizobium meliloti* 1021 *exoY* mutant induces central VB-containing non-nitrogen fixing nodules on wild-type plants (Guan et al., 2013). Detailed cytological analysis of the central VB containing *lin-1* nodules revealed the higher mitotic activity of pericycle and endodermal cells compared to the root cortex, indicating that synchronized mitotic division of cortical, pericyclic, and endodermal layers is necessary for the peripheral VB development (Ting Xiao et al., 2014). In literature, two types of spontaneous nodules or nodule-like structures have been reported. Notably, misregulation of upstream signal transduction pathways can lead to the development of spontaneous nodules with peripheral VB in the absence of bacteria. On the contrary, when downstream transcription factors are overexpressed, nodule-like structures were formed with a central VB (Gleason et al., 2006; Tirichine et al., 2006b; Tirichine et al., 2006a; Tirichine et al., 2007; Soyano et al., 2013; Ried et al., 2014; Saha et al., 2014; Dong et al., 2021; Fu et al., 2022). Immature *M. truncatula* nodules maintain a bilateral symmetry where two vascular traces run parallelly and maintain a connection with the root. In a mature nodule, vascular traces then branch off, resulting in a more intricate network of vasculatures (Guinel, 2009).

Legumes control nodule number via a complex systemic communication known as autoregulation of nodulation (AON). Many of the root-specific AON pathway genes are expressed exclusively in the root phloem cells (Cervantes-Pérez et al., 2022). *M. truncatula Compact Root Architecture* (*CRA2*) encodes a *Leucine-rich repeat receptor-like kinases* (LRR-RLKs). C-terminally Encoded Peptide (CEP) is produced in the root and acts systemically via the MtCRA2 receptor to enhance nodulation and suppress lateral root initiation. Conversely, MtEmbryo Surrounding Region (CLE) peptides also work in a systemic pathway to inhibit nodulation by interacting with SUNN (SUPER NUMERIC NODULES) receptor (Kang et al., 2016). *Too Much Love, TML1,* and *TML2* are negative regulators of nodule initiation and are expressed in roots. CEP-CRA2 promotes microRNA (*miR2111*) production in the shoot and this systemic microRNA promotes the degradation of *TML1/TML2* and initiates nodulation (Tsikou et al., 2018; Gautrat et al., 2020).

*MtbHLH1* is the only transcription factor that has been linked with an unusual nodule VB. To deploy Chimeric REpressor Silencing Technology (CRES-T); *MtbHLH1* was fused to a repression domain and overexpressed in *M. truncatula* to create a dominant negative version of MtbHLH1. *CRES-T-MtbHLH1* nodules showed abnormal branching of peripheral VB. *MtbHLH1* shows expression in the dividing nodule pericycle cells and the VBs of the nodules (Godiard et al., 2011). Here we show, using *M. truncatula Tnt1* insertional mutants of *Mtbhlh1* (renamed as *Nodule Vascular bundle Development 1*) is required for the normal arrangement of vascular strands on the *M. truncatula* nodules periphery. Further, MtbHLH1/NVD1 directly activates a second bHLH group of transcription factor (*NVD2*). *Tnt1* insertional mutants of *NVD2* are also defective in orderly nodule peripheral vasculature development. The stoichiometry of the heterodimer formed by NVD1 and NVD2 is essential for peripheral nodule vasculature development. Auxin and Auxin Responsive transcription Factor (MtARF5) activate *NVD1* and via a feedback loop NVD1/NVD2 control upstream auxin and CK signaling pathways. Our results establish that the NVD1/NVD2 duo works together to control the development of a well-organized peripheral vascular network.

## Results

### *MtbHLH1/NVD1* controls vascular bundle positioning in the nodule periphery

*MtbHLH1* (MtrunA17_Chr3g0132471) is expressed ubiquitously in most tissues, with particularly high levels in roots and several-fold inductions during nodule development (MtExpress)(Carrere et al., 2021). It is one of the earliest transcription factors, beginning to be expressed ∼14 hours post inoculation (hpi), and continuing up to 168 hpi (Supplemental Fig. 1A-B). Noteworthy, after rhizobial inoculation, the first cell division started in the pericyclic layers ∼24 hpi (Ting Xiao et al., 2014). In mature *M. truncatula* nodules, *MtbHLH1* shows the highest expression in the interzone and nitrogen fixation zone, along with lower but still significant expression in the nodule meristem (Supplemental Fig. 1C) (Roux et al., 2014; Carrere et al., 2021). To get a complete understanding of the role of *MtbHLH1* in nodule development we isolated three independent *Tnt1* (*transposable element of Nicotiana tabacum cell type1*) insertional mutants of *MtbHLH1*. Based on the nodule phenotype of mutants in this gene (see below), we renamed this gene as *Nodule Vascular bundle Development 1*. Mutant line NF4649 (*nvd1-1*) has *Tnt1* insertion in the *5’-UTR* of exon 1, NF13338 (*nvd1-2*) has *Tnt1* insertion at 6 bp after start codon, and NF5171 (*nvd1-3*) has *Tnt1* insertion in exon 1 at 59 bp after start codon (Fig. 1A). Plant growth was similar for each of the mutants and wild type (WT) under full nitrogen fertilizer 30 days after sowing (DAS) (Fig. 1B), moreover *nvd1* plants did not exhibit root developmental defects (Supplemental Figure 2A). When grown under the symbiotic condition, none of the mutants show slower plant growth till 21 days post-inoculation (dpi) (Fig. 1B). *M. truncatula* nodules start fixing nitrogen as early as 8 dpi based on the acetylene reduction assay (ARA)(Sinharoy et al., 2013). Hence, we conducted ARA on the *nvd1* and WT plants at three-time points; 10, 15 and 21 dpi. Sporadic differences in ARA could be detected between WT and mutants, but by 21 dpi all three *nvd1* plants showed a significant reduction in ARA (Fig. 1C). In corroboration with the ARA data, reduction in plant fresh weight started appearing only by 30 dpi (Fig. 1B). *nvd1* mutants developed the same number of nodules by 30 dpi as the WT (Fig. 1D and Supplemental Fig. 2B-E). However, closer observation of the macroscopic structure of nodules revealed that *nvd1* nodules were significantly longer than those of the WT, and ∼30 % of nodules of the mutants were distorted compared to the uniform cylindrical structure of WT nodules (Fig. 1D-I and Supplemental Fig. 2B-E). Dissection of distorted *nvd1* nodules contained wavy and partially centralized vasculature (compare Fig. 1J with K-M). Proteins belonging to the same orthogroup were used to generate a phylogenetic tree. Phylogenetic analysis showed that NVD1 belongs to a legume-specific clade (Supplemental Fig. 2F).

**Figure 1.**
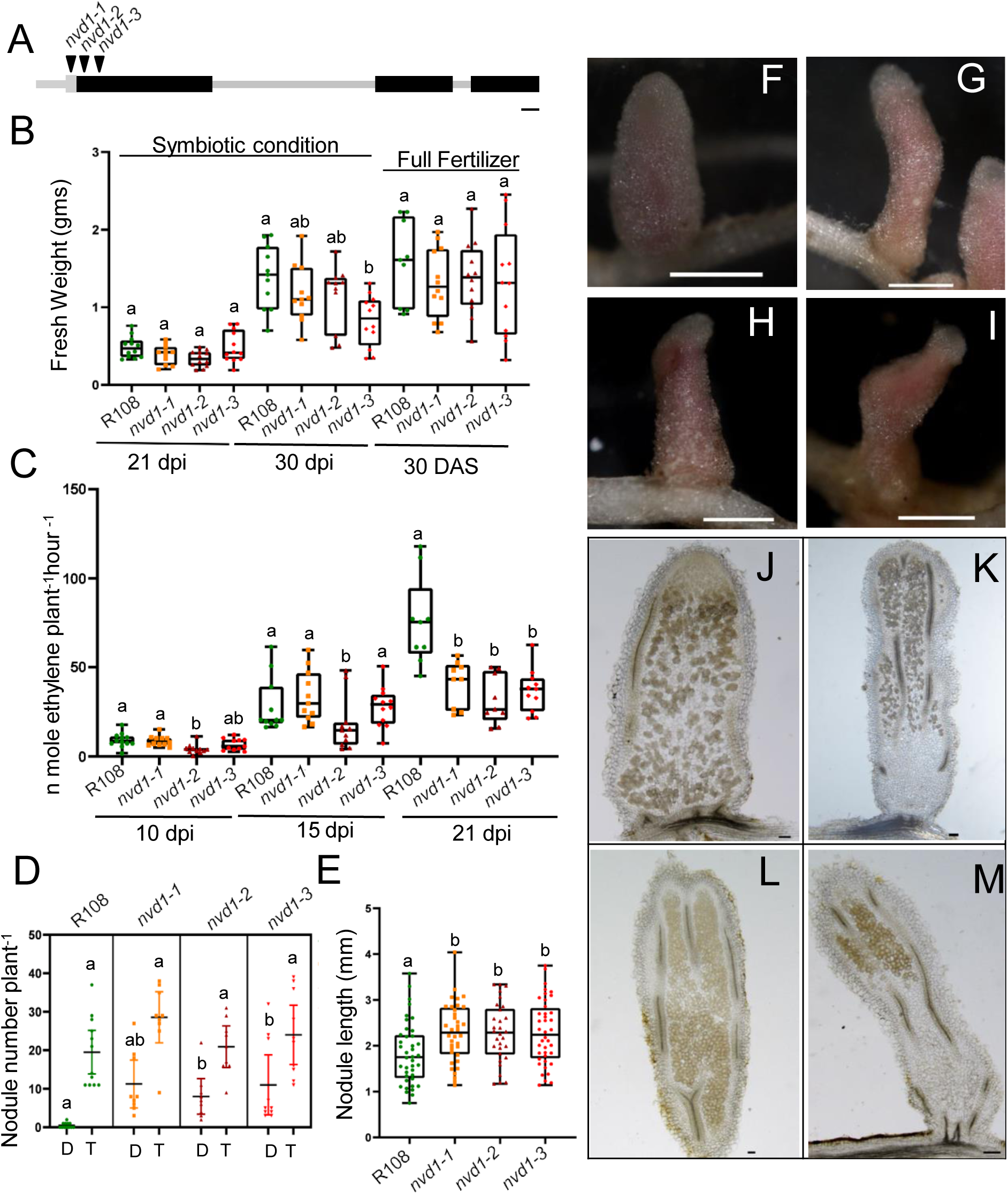
The symbiotic phenotype of *nvd1* mutants. A) Schematic representation of *NVD1* gene model with *Tnt1* insertion positions (arrowheads). Exons are shown in black bars, and introns, promoters, and UTR are shown in gray. Bar = 100 bp B) Fresh weight of wildtype (R108) and *nvd1* plants under symbiotic condition after inoculation with *S. meliloti* 1021 at 21- and 30-days post inoculation (dpi) and non-symbiotic condition by growing plant in presence of full-strength B&D at 30 days after sowing (DAS) (n ≥ 9). C) Acetylene reduction assay (ARA) of R108 and *nvd1* plants at 10, 15, and 21 dpi (n ≥ 9). D) Total nodules and distorted nodule number per plant in R108 and *nvd1.* Distorted nodules are denoted by (D) and the total nodules are denoted by (T) (n ≥ 9). E) R108, *nvd1-1*, *nvd1-2* and *nvd1-3* nodule lengths. Data were collected from 9-12 independent plants. Dots are individual nodule lengths (n ≥ 30) at 30 dpi. The significance of the means was determined within the same time points according to one-way ANOVA and Tukey’s test (P<0.05). (F-M) Macroscopic and microscopic phenotype of *nvd1* nodules. (F-I) Stereo view of R108 and distorted *nvd1* nodules. (A-E) Dots are individual data points. (J-M) cross-section of R108 and *nvd1* nodules. R108 (F, J) *nvd1-1* (G, K) *nvd1-2* (H, L) *nvd1-3* (I, M). Bars = 100 µm (J-L), 150 µm (M), and 1 mm (F-I).

### MtbHLH1/NVD1 activates an HLH transcription factor by directly binding to its promoter

To understand how NVD1 controls NVB development, we performed transcriptome analysis of *nvd1* mutants at 4 dpi (unpublished data). All subsequent experiments were only conducted using *nvd1-2* and *nvd1-3* as both have an insertion in the protein-coding region, hence expected to be more uniform. NVD1 is a transcription activator that binds to a so-called G-box in the promoter of target genes (Godiard et al., 2011). We identified a second bHLH transcription factor (TF), that is expressed at lower levels in the *nvd1* nodules than in the WT. This bHLH-TF has 52 % amino acid sequence similarity with NVD1 but does not have the basic DNA binding part, hence is an HLH protein likely with only protein-protein interaction capacity (Supplemental Fig. 3). We designated *NVD2* (MtrunA17_Chr5g0437991) as the new name for this HLH-TF, due to its significant role in nodule VB development as outlined in the paper. Like NVD1, NVD2 also belongs to a legume-specific clade (Supplemental Fig. 4). *NVD2* expression was significantly decreased in *nvd1* (Fig.2A), suggesting that it may be a direct target of NVD1 TF activity. Unlike *NVD1*, *NVD2* showed feeble expression in roots, but co-expressed with *NVD1* during nodule development and in nodule zones (Supplemental Fig.5 A-B). We further analyzed the NVD2 promoter (*pNVD2*) for G-box elements. About 1 kb upstream of the start codon, *pNVD2* contains six G-box elements with three of these within 185 bp of the start codon (Fig. 2B). Yeast 1 hybrid assays (Y1H) using *pNVD2^185^ ^bp^* showed that NVD1 binds to the *pNVD2^185^ ^bp^*(Fig. 2C). We used 1 kb and 185 bp *pNVD2* for transient transactivation assay using *Nicotiana benthamiana* leaves, following co-infiltration of *p35S::NVD1* (Fig. 2D-E). The use of both 1-kb and 185-bp promoters led to the induction of *NVD2* expression (Fig. 2D-E). These results establish that *NVD2* is a direct target of NVD1.

**Figure 2.**
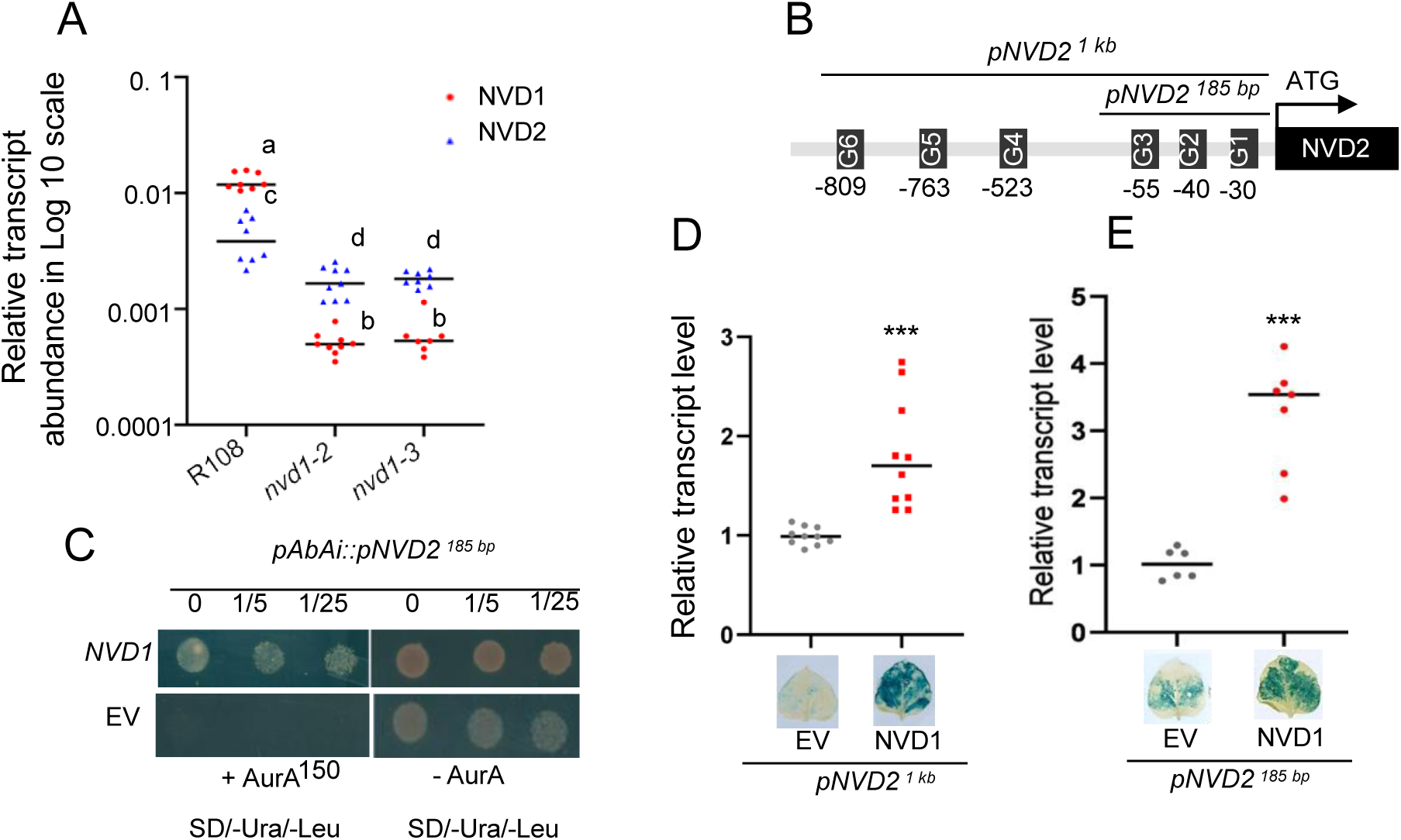
NVD1 directly activates the expression of *NVD2*. A) Relative transcript abundance of *NVD1* and *NVD2* in R108, *nvd1-2,* and *nvd1-3* at 15 dpi. Dots are individual data points from two biological and three technical replicates. B) Schematic diagram of *NVD2* promoter. The G-boxes (CANNTG) were denoted along the *pNVD2*. *pNVD2^1kb^* and *pNVD2^185bp^* were marked, used for Y1H and tobacco transactivation assays in (C-E). *pNVD2*^185bp^ contains three and *pNVD2^1kb^* contains six G-box elements. (C) Interaction of NVD1 with *pNVD2*. *pNVD2*^185bp^ was used to create the Y1HGold bait reporter strain. *NVD1* was cloned in the *pGADT7* vector and used as prey. Yeast cells containing either *pGADT7 vector or pGADT7::NVD1* were grown without Ura and Leu containing medium in the presence or absence of Aureobasidin A (150 ng). Yeast cells were platted in 0, 1/5, and 1/25 dilutions. (D-E) Transactivation assay using different lengths of *pNVD2::GUS* construct co-infiltrated with NVD1 in *N. benthamiana* leaves. (D) *pNVD2^1kb^* and (E) *pNVD2^185 bp^*. Relative transcript abundance of *GUS* in transformed leaves containing respective constructs, *nptII* used as a housekeeping gene. Representative GUS-stained leaf images are given below. Dots are individual data points from three biological and two or more technical replicates and each replicate contains a pool of four leaves.

### NVD1 and NVD2 form a heterodimer and control nodule vascular bundle development

bHLH TFs have a ∼60 amino acid signature domain, with an N terminal part consisting of ∼15 basic amino acids, followed by a hydrophobic helix-loop-helix region. Structural studies have demonstrated that protein-protein interaction between the HLH regions of two separate polypeptides leads to the formation of a functional DNA binding fork (Toledo-Ortiz et al., 2003). Hence, we modeled the homodimeric structure of NVD1 and the heterodimeric structure of NVD1-NVD2 (Fig. 3A-B) using alpha-fold (Jumper et al., 2021). Structural analysis indicated that the NVD1-NVD2 heterodimer forms a distorted DNA binding fork (Fig. 3B). Next, we performed a luciferase complementation assay in *N. benthamiana* leaves involving transient transfection with Agrobacterium to test the heterodimer formation capacity of NVD1 and NVD2. At 36-hour post infiltration, NVD1 and NVD2 showed strong heterodimer formation as evident from luminescence (Fig. 3C). NVD1 activates *NVD2* during nodule development (Fig. 2C-E), and NVD2 can heterodimerize with NVD1 possible inhibiting its DNA binding. As evident from the alpha-fold predicted model NVD1 and NVD2 heterodimer might not be able to activate transcription. We hypothesized that, when NVD2 is expressed under the early nodule-inducing ENOD12 promoter (*pENOD12*), it can form a heterodimer with NVD1 and consequently sequester it, possibly inhibiting its DNA binding. In such an experiment the stoichiometry of the homo- and hetero-dimer will be distorted. To test this hypothesis, we generated transgenic nodules expressing *pENOD12::NVD2* construct and the ectopic expression of *NVD2* was validated by qRT-PCR (Fig. 3D). Interestingly, ectopic expression of *NVD2* led to the downregulation of *NVD1*. *pENOD12::NVD2* transformed nodules contained wavy and partially central nodule vasculature like *nvd1* (compare Fig. 3E and F). To determine the reason for reduced *NVD1* in transgenic *pENOD12::NVD2* nodules, we analyze the 2 kb upstream region of the *NVD1* promoter (*pNVD1*). The use of 2 kb *pNVD1* led to a specific expression pattern of *NVD1* in *M. truncatula* nodules, as confirmed by in situ hybridization (Godiard et al., 2011). We detected five G*-boxes* in *pNVD1^2kb^* and tested whether NVD1 can activate its own expression. Transactivation assay in *N. benthamiana* leaves showed that co-infiltration of *p35S::NVD1* can activate *pNVD1^2kb^::GUS* expression (Fig. 3G). Thus, NVD1 can activate its own expression. To biochemically validate NVD2 medicated NVD1 regulation, we co-infiltration *pNVD1^2kb^* and *pNVD2^1kb^*along with either *p35S::NVD1* or both *p35S::NVD1* and *p35S::NVD2*. Simultaneously introducing NVD1 and NVD2 leads to the suppression of NVD1-mediated activation of both *NVD1* and *NVD2*. Therefore, during nodule development, NVD2 exerts control over NVD1 through two mechanisms, i) repression of NVD1 expression at the transcriptional level, as NVD1 activates its own expression in a feed-forward loop (Fig. 3D and G) ii) NVD2 binding to NVD1 and forming a non-functional heterodimer, which regulates NVD1’s activity towards its downstream targets.

**Figure 3.**
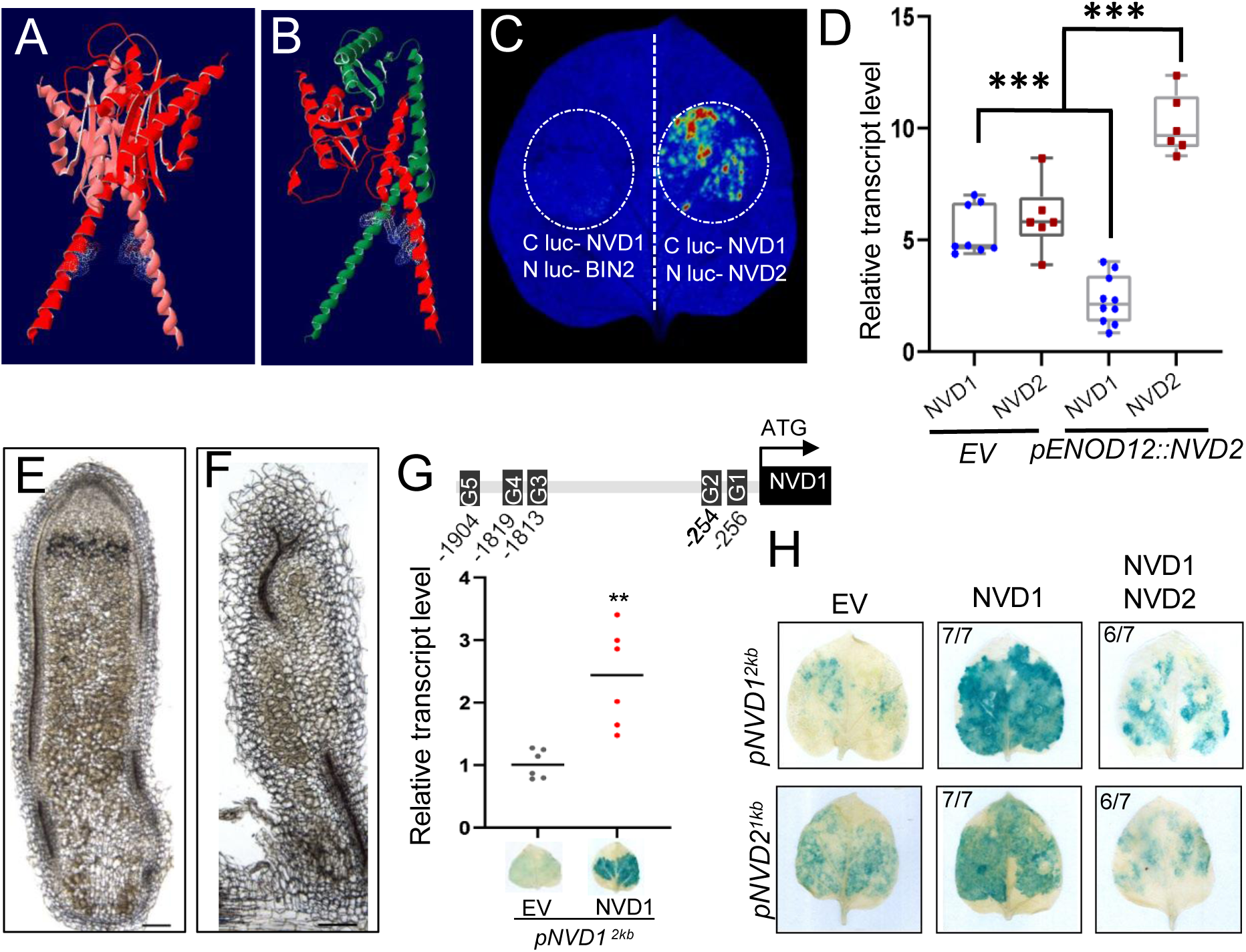
NVD1 and NVD2 form a heterodimer and regulate nodule vascular bundle positioning. (A-B) Alpha fold (Mirdita et al., 2022) is used to predict NVD1-NVD1 homodimer (A), NVD1-NVD2 heterodimer (B). Visualization of the structure was done using Swiss-PdbViewer v 4.1.1 (Guex and Peitsch, 1997). NVD1 polypeptide chains are depicted in red and light red in (A) and NVD1 and NVD2 polypeptides are depicted as red and green respectively in (B). The electron cloud of the invariant amino acids that are required for DNA binding is shown (see Supplemental Fig. 3). (C) Split luciferase (LUC) complementation imaging assays between NVD1 and NVD2. cLUC (C terminus of LUC) was tagged with *NVD1* and infiltrated in *N. benthamiana* leaves with *nLUC-NVD2* or *nLUC-AtBIN2* (AT4G18710.1) (negative control). (D-F) Ectopic expression of *NVD2* using *pKGW-RedRoot-proENOD12-eGFP* (Sinharoy et al., 2016) in R108 through hairy root transformation. (D) Relative transcript abundance of *NVD1* and *NVD2* in empty vector (EV) control plants and ectopically expressed *pENOD12::NVD2* hairy root transformed nodule at 15 dpi. Dots are individual data points from three biological and two or more technical replicates. Student t-test was used to generate the statistical significance. ***, P<0.001. Cross section of the transformed nodule with EV control (E), and ectopically expressed *pENOD12::NVD2* nodules (F) at 30 dpi. (G) Transactivation assay of *pNVD1*::GUS construct co-infiltrated with *p35S::NVD1* in *N. benthamiana* leaves. (H) Co-infiltration of NVD1 with NVD2 suppress the transactivation capacity of NVD1. Transactivation assay of *pNVD1^2kb^::GUS* and *pNVD2^1kb^::GUS* construct co-infiltrated with *p35S::NVD1* or *p35S::NVD1* and *p35S::NVD2* in *N. benthamiana* leaves. Representative image of leaves was given. Number of positive tested leaves/total number of leaves analyzed is given in (H). Scale Bar = 100 µm.

### The presence of NVD2 is crucial for peripheral nodule vascular strand development

Ectopic expression of *NVD2* reduces the functional DNA-binding units of the NVD1 homodimer recapitulating the *nvd1* phenotype. To understand at the genetic level how NVD2 controls peripheral NVBs development, we isolated two independent *Tnt1* mutant lines of *NVD2*. NF7653 (*nvd2-1*) contains a *Tnt1* insertion in exon 3, at 347 bp after the start codon, while; NF19150 (*nvd2-2*) contains a *Tnt1* insertion in exon 4 in the *3’-UTR* (Fig. 4A). *NVD2* expression was significantly reduced in *nvd2-2* nodules, but not in *nvd2-1* compared to the WT, R108 (Fig. 4B). Previous studies have found that exonic *Tnt1* insertions do not necessarily lead to a decrease in transcript level (Roy et al., 2017; Liu et al., 2019). Phenotyping revealed *nvd2-1* and *nvd2-2* have similar defects (Fig. 4C-I). At an earlier stage of nodule development up to 30 dpi, no significant reduction in ARA was detected in *nvd2* nodules compared to the WT, R108 (data not shown). However, at 45 dpi, ARA was significantly lower in *nvd2-1* and *nvd2-2* plants (Fig. 4C). Similarly, *nvd2* mutants exhibited significantly lower plant fresh weight than the WT at 45 dpi (Fig. 4D). Like *nvd1*, nodule number was unaffected in *nvd2* mutants (Fig. 4E). Taken together, these results indicate that the *NVD2* mutation produces a less potent effect on nitrogen fixation and plant growth than *nvd1*. Further, the macroscopic deformation of nodules observed in *nvd1,* was absent in *nvd2* nodules (Fig. 4F). Nonetheless, dissection of *nvd2* nodules showed similar vasculature defects as *nvd1*, with chaotic, partially centralized vascular bundles (compare Fig. 4G with H and I). The deformation of vascular bundles in *nvd2* nodules was usually more prominent towards the nodule apex, in contrast to *nvd1* nodules where deformed vascular bundles were noticed throughout the nodule (Compare Fig. 1J-M and Fig. 4G-I). This difference in the degree of vascular bundle re-positioning may account for the absence of a macroscopic defect in *nvd2* nodules. We summarized that the absence of *NVD2* results in an abundance of functional NVD1 homodimers, with *nvd2* nodules phenocopying *NVD1* overexpression. Our data indicate that the correct stoichiometry of NVD1 and NVD2 is crucial for maintaining peripheral nodule vasculature.

**Figure 4.**
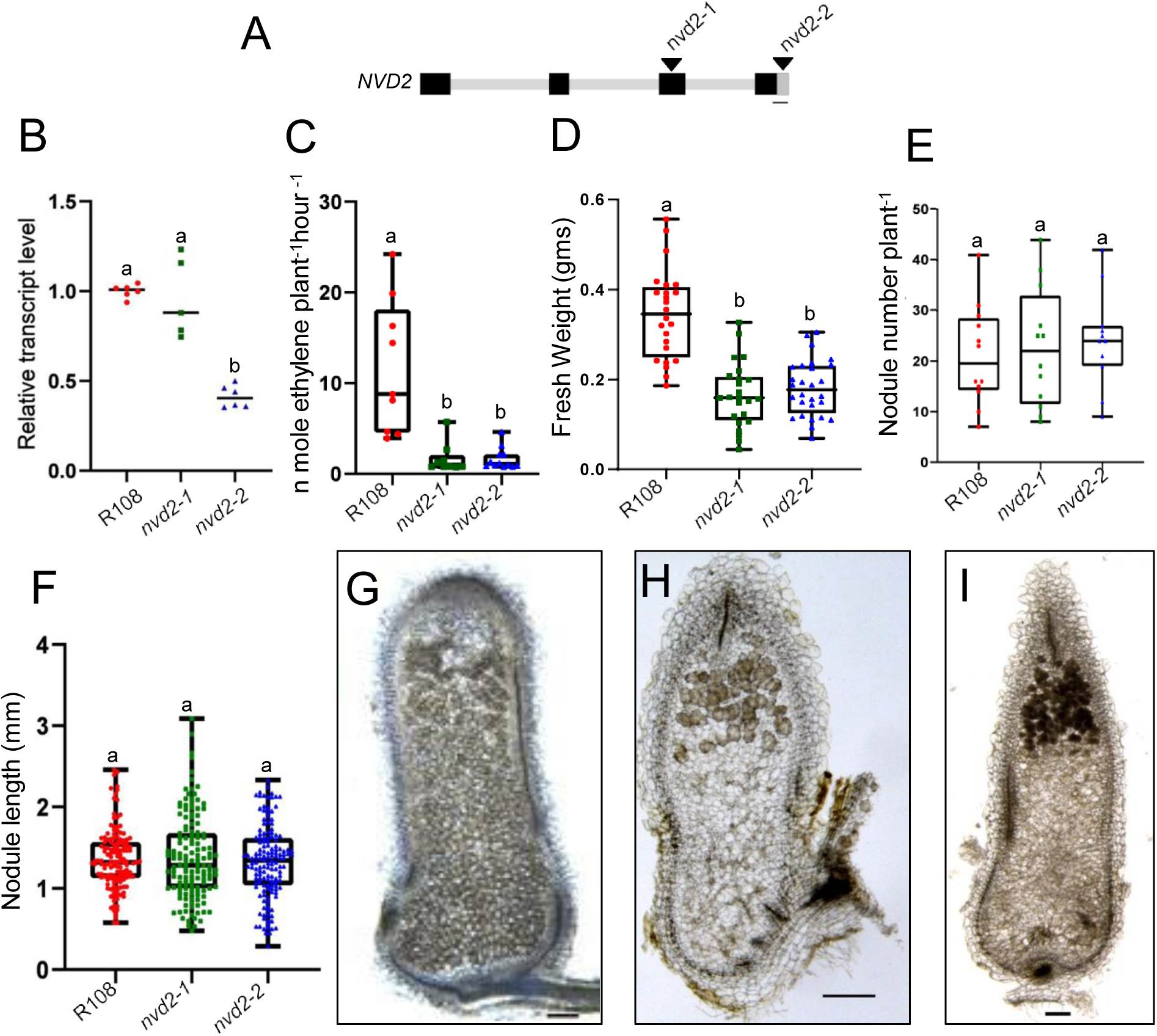
The symbiotic phenotype of *nvd2* mutants. A) Schematic representation of *NVD2* gene model with *Tnt1* insertion positions (arrowheads). Exons are shown in black bars, and introns and UTR are shown in gray. Bar = 100 bp. B) Relative transcript abundance of *NVD2* in R108, *nvd2-1,* and *nvd2-2* at 15 dpi nodules. Dots are individual data points from two biological and three or more technical replicates. (C-D) Evaluation of the performance of the plants under symbiotic conditions after inoculation with *S. meliloti 1021* at 45 dpi. ARA (C), fresh weight (D) of R108 and *nvd2* plants (n ≥ 9). Dots represent data collected from individual plants. (E) number of nodules in R108, *nvd2* plant. Dots are individual data points (n ≥ 9). (E) Length of R108 and *nvd2* nodules in mm at 30 dpi. Dots are individual data points from 11 independent plants (n ≥ 138) and (G-I) Cross-section of R108 (G), *nvd2-1* (H) and *nvd2-2* (I) nodules at 30 dpi. Scale bar = 100 µm.

### *MtNVD2* is expressed in the nodule primordium, developing nodule vascular bundles and nodule vascular meristem

To determine the spatiotemporal expression pattern of *NVD2,* we used *pNVD2^1kb^::eGFP:GUS* and *pNVD2^185bp^::eGFP:GUS* constructs expressed in *M. truncatula* R108 ecotype using rhizogenes-mediated hairy root transformation. Transgenic *pNVD2^1kb^::eGFP:GUS* roots showed *NVD2* promoter activity in the root apex and stele (Fig. 5A). In developing nodules, at stages III-IV, *NVD2* was expressed strongly in the inner cortical layers (C4-C5), endodermis, pericycle, and root stele, and its expression was maintained throughout the nodule primordium and root stele at stage V (Fig. 5C). The shorter *pNVD2^185bp^::eGFP:GUS* construct resulted in GUS expression mainly in the nodule vasculature and root stele in a mature *M. truncatula* nodule at 20 dpi (Fig. 5D) similar to a longer 1 kb promoter (Supplemental Fig. 5C). A short period of GUS staining of nodules of *pNVD2^1kb^::eGFP:GUS* hairy roots revealed *NVD2* expression restricted in the NVMs at 20 dpi but not present in the NCM (Fig. 5E). In nodule cross sections, we detected strong expression of *NVD2* in the NVB including nodule vascular vessel, nodule vascular endodermis (NVE) and nodule vascular pericycle (NVP), but not in nodule parenchyma (NP) and nodule endodermis (NE) (Fig. 5F). To understand cellular localization of NVD2, we created *pNVD2^1kb^::NVD2:eGFP* construct and generated composite plants with transgenic roots. In a nodule longitudinal section, NVD2:GFP fusion protein is localized in the nucleus of the vascular bundle cells (Fig. 5G). Whereas in a nodule cross-section, NVD2:GFP fusion localized specifically in the nucleus of the NVE (Fig. 5H). We determined that NVD2 is expressed more widely in nodule primordium during the initial stages of nodule growth, but its expression is limited to the NVB at a later stage. We detected specific localization of NVD2 in the NVE in a mature *M. truncatula* nodule.

**Figure 5.**
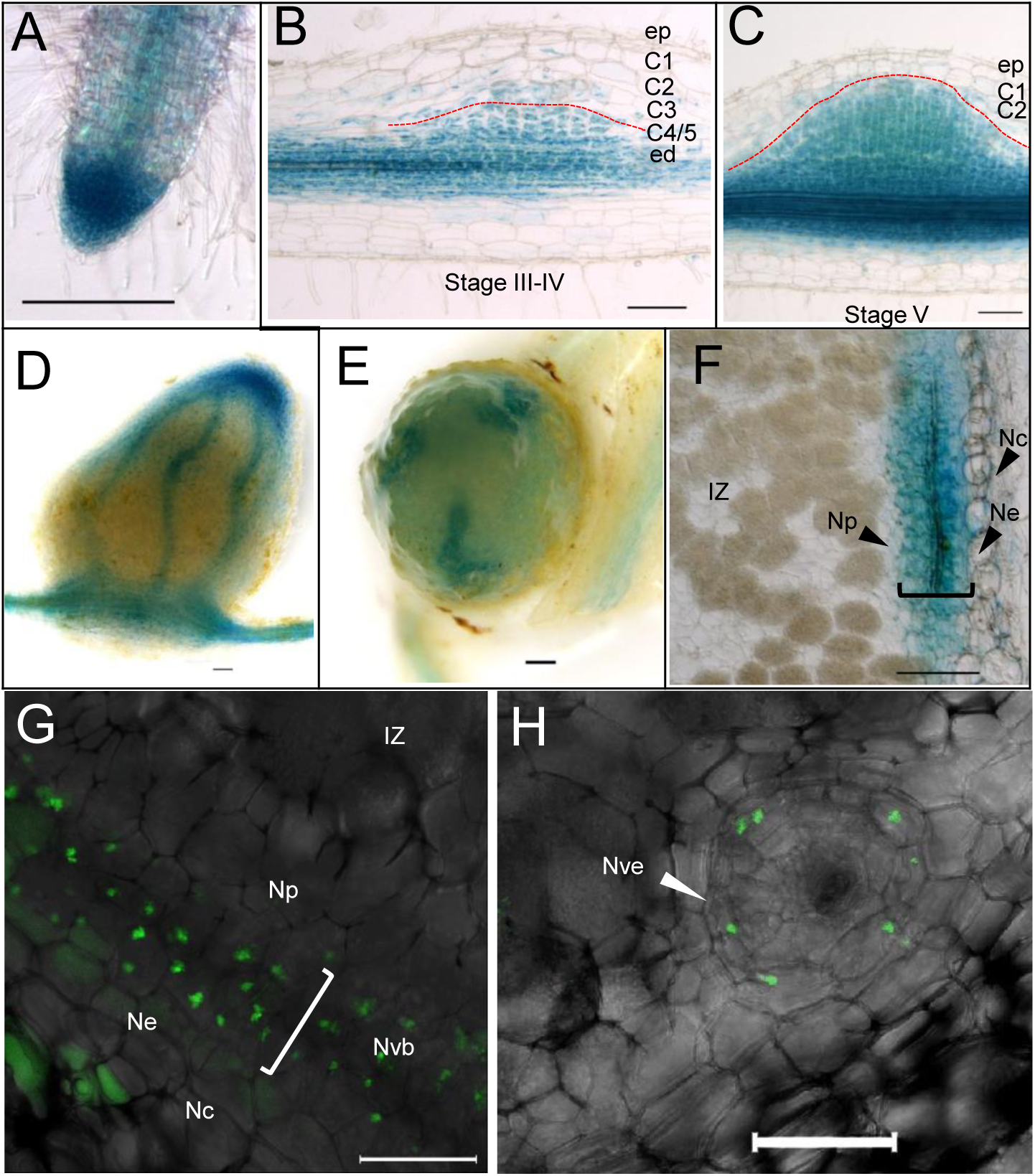
NVD2 expresses in nodule vascular bundles and vascular meristems, and NVD2-GFP localizes to the nucleus of nodule vascular endodermis. (A-C) Representative image showing GUS activity using pNVD21kb::GUS transformed transgenic roots, in apical root meristem and root vascular bundle (A), dividing root cortical cell with GUS activity at nodule developmental stage (III-IV) (B), Nodule primordium at nodule developmental stage (V) (C). Red lines demark the border of C4/5 from C3 in (B) and C1/2 from nodule primordium (C). Cell layers and stages are marked according to the M. truncatula fate map (Ting Xiao et al., 2014). (D) Representative image of pNVD2185bp::GUS transformed 20 dpi nodule shows GUS activity in nodule vascular bundles (Nvb) (see Supplemental Figure 5C for pNVD21kb:: GUS at 20 dpi). Staining was performed for 3 hours for (A-D) (E) Representative image of pNVD21kb::GUS transformed nodule shows GUS activity in nodule meristem, particularly in NVM after 1 hour of staining. (F) Section of 20 dpi nodule after GUS staining. pNVD21kb activity in nodule endodermis (Ne) and Nvb; (G-H) Maximum intensity projection of the merged confocal images showing localization of NVD2-GFP fusion protein in nodule sections after transforming with pNVD21kb::NVD2:GFP. Representative image of a longitudinal nodule section showing NVD2-GFP in the nucleus of Vb (G), transverse section showing NVD2-GFP in the nodule vascular endodermis (Nve) (H). Arrow indicates Nve. Ed, endodermis; ep, epidermis; C, cortex; Np, nodule parenchyma; Nc, nodule cortex; IZ, Infection zone; NVM, Nodule vascular meristem. Bars=100µM (A,F); 50 µM (B-E, G, and H).

### Comparable changes in the transcriptomes of *nvd1* and *nvd2* nodules

To understand how NVD1 and NVD2 control NVB development, we performed transcriptome analyses of *nvd1* and *nvd2* nodules at 15 dpi and compared gene expression with wild type (R108). We detected 2278 (1267 downregulated and 1011 upregulated) and 5407 (2481 downregulated and 2925 upregulated) differentially expressed genes (DEGs) in *nvd1* and *nvd2* nodules respectively (Fig. 6A). Out of these DEGs, 1672 were found to be common to both mutants (Fig. 6B). Among the common DEGs, there were 722 upregulated genes and 946 downregulated genes that showed the same direction of deregulation in both mutants (Supplemental Figure 6A-D). We compared common DEGs of *nvd1* and *nvd2* (together *nvd1* and *nvd2* will be referred to as *nvd* from now onward) with *M. truncatula* nodule zonation data (Roux et al., 2014) and detected a higher proportion of DEGs amongst nodule meristem and invasion zone specific genes (Fig. 6C). This suggests that *nvd* nodules are largely affected in terms of nodule meristem development/maintenance. Noteworthy, both *NVD1* and *NVD2* displayed expression at the nodule meristem (Supplemental Figure. 1C and 5B).

**Figure 6.**
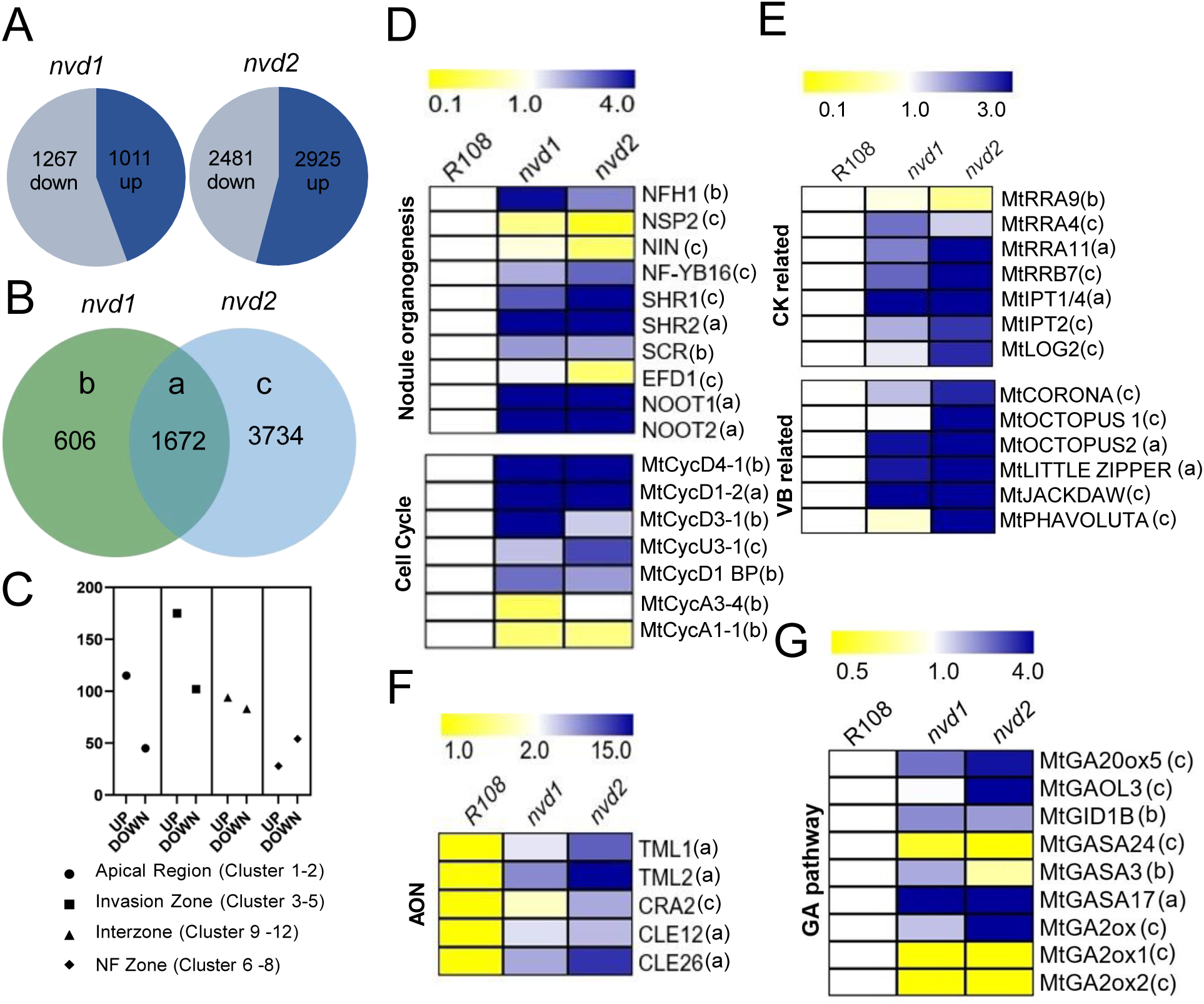
Transcriptome analysis of *nvd1* and *nvd2* during nodule development. A) Summarization of up and down-regulated genes in *nvd1* and *nvd2* nodules at 15 dpi compared with wild type R108. B) Venn diagram showing the number of differentially expressed genes (DEGs) in *nvd1* and *nvd2* nodules. Overlapping DEGs shared by both *nvd1* and *nvd2* labeled as (a), DEG specific to *nvd1* nodules labeled as (b), and DEGs specific to *nvd2* labeled as (c). C) Comparison of overlapping DEGs with the laser microdissection data of nodule zones (Roux et al., 2014) (see Supplemental table 1). D-G) Heatmap showing the relative transcript abundance of selected genes in R108, *nvd1,* and *nvd2* related to nodule organogenesis and cell cycle (D), Cytokinin CK) related and vascular bundle (VB) related (E), AON (autoregulation of nodulation) related (F) and GA (Gibberellic Acid pathway) related (G). The scale bar depicts the fold-change for D-G.

We categorized the DEGs of *nvd1* and *nvd2* nodules mainly in four clusters. Among them, the majority of genes were upregulated in the mutants compared to the WT (Fig. 6D-G). The first cluster consists of development-related genes including *MtSHR1/2*, *MtSCR*, *MtNOOT1/2,* and *MtNF-YB16* which were 2-4-fold upregulated, and symbiosis-specific transcription factors *NIN*, *NSP2,* and *EFD1,* which were 2-5-fold downregulated (Fig. 6D). *SHR2*, *NOOT1,* and *NOOT2* are common DEGs, whereas rest of the genes were DE in either *nvd1* or *nvd2* nodules (Fig. 6D). We also detected upregulation of D-type cyclins, cyclin D binding proteins, and plant-specific cyclin U. In contrast, rhizobium-induced cyclins get down-regulated (Fig. 6D). Cyclins are mainly upregulated in *nvd1* nodules except *MtCycD1-2* which is upregulated in *nvd* nodules and *Cyclin U*, which is an *nvd2* specific upregulation. Noteworthy, in Arabidopsis AtSHR activates D-type cyclins and promotes cell division to separate endodermis and cortical cell layers (Shaar-Moshe and Brady, 2022). GmSHR4/5 activate *cyclin D*s during nodule development (Wang et al., 2022). We also detected upregulation of *MtNFH1* in *nvd1* nodules. Notably, disruption of NF balance leads to the formation of multiple nodule meristems (Cai et al., 2018).

In the second cluster, we have detected the induction of CK biosynthesis-related genes. *MtIPT1/4* is a common upregulated gene, while *MtIPT2* and *MtLOG2* are specifically upregulated in *nvd2* nodules. Regarding RRs, three *type A* and one *type B* are differentially regulated in *nvd* nodules. Among them, *MtRR9* is downregulated in *nvd1* nodules, while all other RRs are upregulated, and *MtRR11* is the only *RR* common among *nvd* nodules. Notably, NF treatment activates *MtRR11* and *MtRR9* within 3 hours (Op den Camp et al., 2011). Among 190 auxin/CK pathway-related genes that are present in the *M. truncatula* genome, 69 are DE in *nvd* nodules (Supplemental Table 2). Among them seven *ARFs*, eight *Aux/IAA*, two *PINs,* and four *YUCCAs* are present, all indicated towards perturbation in the auxin signaling. Together this data suggested that the balance of auxin and CK signaling was disrupted in the *nvd* nodules. Furthermore, we also detected, *M. truncatula* orthologues of Arabidopsis genes that are required for root vascular bundle development such as *CORONA* (Green et al., 2005), *OCTOPUS1/2* (Anne et al., 2015), *Little-Zipper*, *Phavoluta* (Wenkel et al., 2007), and *Jackdaw* (Fig. 6E). Among these vascular bundle-related genes, most were upregulated only in *nvd2* nodules, while *MtOCTOPUS2* and *MtLittle-Zipper* were commonly upregulated genes (Fig.6E).

In cluster three, we detected an upregulation of AON-related genes, namely *CRA2*, *TML1/2*, *CLE12*, *CLE26* (Fig. 6F). Except for *CRA2*, these genes were upregulated in *nvd* nodule (Fig. 6F). However, despite the disruption in the AON pathway, we did not observe any changes in nodule numbers in *nvd* plants (Fig. 1D and 4E). Interestingly, despite perturbation in the AON pathway nodule number is not changed in *nvd* mutants (Fig. 1D and Fig. 4E). Moving on to cluster four, we observed upregulation of the gibberellic acid (GA) biosynthesis, along with downregulation of GA catabolism-related genes, suggesting a disturbance (most likely accumulation) of GA signaling inside the *nvd* nodules. Active GA stimulates *GA-stimulated Arabidopsis* (*GASA*) orthologs. *MtGASA3, 17,* and *24 which* were DE in *nvd* nodules (Fig. 6G). The majority of DEGs belonging to this cluster were specific to *nvd2*. In particular, *GASA3*, *GASA17,* and *GID-1B* showed DE in either *nvd* nodules or preferentially in *nvd1* nodules (Fig. 6G).

The majority of the DEGs in *the nvd* transcriptome were de-regulated in the same way in both mutants (Supplemental Figure 6A-D). Noteworthy, in *nvd2* nodules, 21 *bHLH-TFs* were upregulated. Among the 21 upregulated *bHLH-TFs*, 7 lack a basic DNA binding domain (Supplemental Figure 6E-F), suggesting a potential feedback regulation among the bHLH family. NVD2 prevents NVD1-mediated gene activation by sequestering NVD1. Hence, we predicted that the potential direct target of NVD1 will be downregulated in *nvd1* nodules and will be upregulated in *nvd2* nodules. In the *nvd* transcriptome, we obtained only 4 genes that satisfy the above-mentioned criteria (Supplemental Fig. 6A and Supplemental Table 1). We hypothesized that the reason behind this apparent anomaly is the DE of a large number of *bHLH-TFs* in *nvd2* nodules.

Though *nvd2* had a milder phenotype, we detected more DEGs in the *nvd2* nodules. Among genes that were specifically upregulated in *nvd2* nodules, a cluster of defense-related genes was present. *Regulator of Symbiosome Differentiation* (*RSD*)(Sinharoy et al., 2013), *Nodules with Activated Defense 1* (*NAD1*)(Wang et al., 2016) were upregulated in *nvd2* nodules along with senescence-related and pathogenesis-related genes (Supplemental Fig. 6G). Numerous NCRs were either up-or down-regulated in one or both mutants. To be specific, 68 *NCRs* were upregulated in *nvd2* nodules (Supplemental Figure 6H). Defense response gets activated in *rsd1,* and *nad1* nodules and leads to the degradation of bacteroids. Again, disruption in the balance of *NCR* genes also promotes the degradation of bacteroids. As the balance among *RSD*, *NAD1,* and *NCRs’* got disturbed, we tested the viability of bacteria inside the *nvd* nodules. Live/dead staining showed that despite this perturbation, rhizobial viability remains non-compromised inside the *nvd* nodules at 15 dpi (Supplemental Fig. 7A-F).

### *NVD1/2* works downstream of auxin signaling and controls auxin/cytokinin balance in nodules via a feedback loop

*MtbHLH1/NVD1* is induced by auxin treatment (Godiard et al., 2011). Detailed analysis of *NVD1* and *NVD2* promoters identified 17 and 12 auxin-responsive elements in *pNVD1^2kb^* and *pNVD2^2kb^* respectively (Fig. 7A). We applied auxin to R108 roots, which induced *NVD1* and *NVD2* expression by 6 hours (Fig. 7B). Auxin responsiveness was higher in the case of *NVD1*. To explore the relationship between auxin and NVD1/2, we sought and identified seven DE *ARF* genes in *nvd1* and *nvd2* mutants, four of which were induced in both mutants compared to the WT (Fig. 7C). *MtrunA17_Chr1g0156321* (*MtARF5*) and *MtrunA17_Chr7g0239061* (*MtARF9*) showed high co-expression (>0.8) with both *NVD1* and *NVD2* according to nodule developmental time series (Schiessl et al., 2019). Further, according to LCM-nodule transcriptome data, *MtARF5*, *NVD1,* and *NVD2* are expressed preferentially in nodule meristem (Roux et al., 2014). The ortholog of MtARF5, AtARF5 in Arabidopsis (also known as MONOPTEROS) is a crucial regulator for root embryogenesis, (Schlereth et al., 2010) and has been characterized as a transcription activator (Tiwari et al., 2003). Hence, we tested if MtARF5 can activate *NVD1* and *NVD2* expression using a transactivation assay in *N. benthamiana* leaves. In this system, both *NVD1* and *NVD2* got activated by MtARF5 (Fig. 7E-F). In light of the fact that NVD1 also induced *NVD2* expression (Fig. 2B-E), we tested whether NVD1 mediates auxin induction of *NVD2*. Auxin treatment of *nvd1* roots did not induce *NVD2* expression even after 12 hours (Fig. 7G). Indicating that NVD1 is necessary for the auxin-mediated induction of *NVD2*.

**Figure 7.**
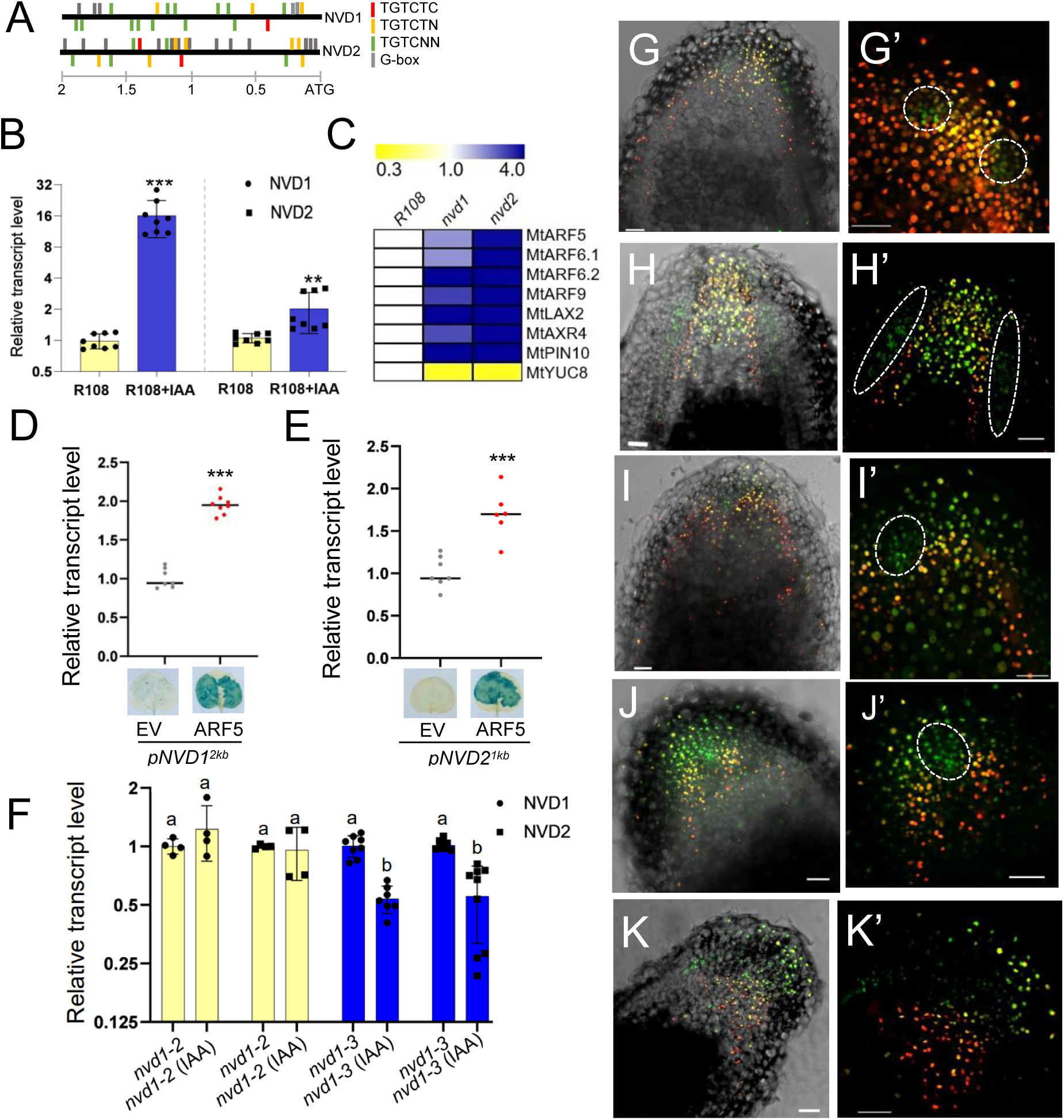
Auxin induces *NVD1* via ARF5 and NVD1-NVD2 controls the auxin distribution pattern at the nodule apex. A) Schematic representation of *pNVD1^2kb^* and *pNVD2^2kb^*. *pNVD1^2kb^* contains seventeen auxin-responsive elements (AuxRE) along with five G-box while *pNVD2^2kb^* contains twelve AuxRE with six G-box. AuxRE is represented on plus (upper side) and minus (lower side) strands. Different AuxRE sequences TGTCTC (Red), TGTCNN (Green), and TGTCTN (Yellow) are represented on the respective promoters. The grey bar represents G-box. B) Relative transcript level of *NVD1* and *NVD2* after auxin treatment. 10 µM auxin (IAA) treatment was given for 6 hours to *M. truncatula* roots. C) Heatmap showing the expression pattern of genes related to auxin signaling and belonging to category (a) in Figure 6B. D) Relative transcript level of *GUS* in *pNVD1^2kb^* or *pNVD1^2kb^* along with *35S::MtARF5* transformed leaves. Representative GUS-stained leaves were shown below. E) Relative transcript level of *GUS* in *pNVD2^1kb^* with/without co-infiltrated *p35::MtARF5.* The representative image of GUS-stained *N. benthamiana* leaves was given in the lower panel. F) Relative transcript level of *NVD1* and *NVD2* after auxin treatment. 20 µM auxin (IAA) treatment was given for 12 hours to *M. truncatula* roots. The significance of the means was calculated using one-way ANOVA and Tukey’s test (P<0.05). G-K’) longitudinal section of nodules transformed with *2COL* vector, showing auxin and cytokinin distribution pattern. G-G’, R108, H-H’, *nvd1-2*, I-I’, *nvd1-3,* J-J’, *nvd2-1,* K-K’, *nvd2-2*. Auxin maxima are shown by GFP-NLS (green) and cytokinin maxima by td-Tomato-NLS (red) in R108, *nvd1* and *nvd2* transformed nodules. A dashed enclosure demarcated the zone of auxin maxima. Statistics were calculated using a student t-test. ***, P<0.001; **, P<0.005 in B, D and E. Scale bar = 50 μm.

A variety of auxin signaling-related genes were aberrantly expressed in the *nvd* nodules, including auxin influx and efflux carrier proteins *MtLAX2* and *MtPIN10* and the putative regulator, MtAXR4 which were induced and MtYUC8 which was expressed at lower levels than WT (Supplemental Table 2; Fig. 7D). Further, CK biosynthesis and response regulators (RR) were also differentially expressed in *nvd* nodules (Fig. 6E), indicating an alteration in the auxin and CK signaling inside *nvd* nodules. Given that interactions between CK and auxin signaling are crucial for plant vascular bundle development (De Rybel et al., 2016) we evaluated nuclear auxin and CK signal outputs in the *nvd* mutants and wildtype (R108), using a construct that contains synthetic auxin (*pDR5*-) and CK (*pTCSn*-) responsive promoter-driven *GFP-NLS* and *tdTomato-NLS* expression, respectively (abbreviated as 2-COL; (Fisher et al., 2018). *2-COL* has been used successfully to monitor auxin and CK responses simultaneously during nodule development (Fisher et al., 2018; Bhattacharjee et al., 2022). We obtained auxin maxima in NVMs and high CK levels in the NCM of WT nodules (Fig. 7G-G’). In the WT nodule, two NVMs are located symmetrically in the periphery of the NCM at the nodule apex (Fig. 7G-G’). Using the auxin maxima as a guide, NVMs were identified in the mutants. In *nvd1-2* nodule NVMs were broader and situated below the nodule apex compared to the WT (Fig. 7H-H’), whereas in *nvd1-3* nodules only one NVM is present in the nodule apex and NCM identity is probably lost (Fig. 7I-I’). Similarly, in *nvd2* nodules, the symmetrical distribution pattern of NVMs was lost. In *nvd2-1* nodule, a broader auxin maximum at the nodule apex indicated that the nodule meristem lost its NCM identity and retained only NVM identity (Fig. 7J-J’). In *nvd2-2* nodules based on hormonal distribution it was challenging to differentiate NCM or NVM (Fig. 7H-H’). Noteworthy, the hormonal distribution pattern in the *nvd1* and *nvd2* nodules displays irregularities that vary from nodule to nodule within a single mutant type. We have identified several distinct patterns of aberrant hormonal distribution, which we have presented here (Fig H-K). Taken together, these data suggest that the mutant nodules lost their characteristic meristem identity, which we believed to be the primary reason behind the abnormal development of peripheral vascular bundles in *nvd* nodules.

## Discussion

In this study, we showed that *MtbHLH1*/*NVD1* and an interacting HLH protein lacking a DNA binding domain, NVD2, control the positioning and patterning of the peripheral vascular bundle in *M. truncatula* nodules. To summarize, we found that: (i) NVD1 induces expression of *NVD2;* (ii) NVD1 and NVD2 can form a heterodimer; (iii) NVD2 controls NVD1 transactivation capacity by sequestering NVD1; (iv) ectopic expression of *NVD2*, recapitulated the *nvd1* mutant phenotype; (v) *nvd1*, *nvd2* nodules show defective vascular bundle patterning; (vi) *NVD1/2* works downstream of auxin and MtARF5 (vi) expression of auxin and CK signaling pathway genes and auxin/CK distribution pattern are disrupted in *nvd* nodules. These data provide insight into the evolution of peripheral vascular development in legume nodules, as opposed to the central vascular development of roots, as discussed below.

*AtARF5/MONOPTEROS* works downstream of auxin signaling and is a crucial determinant of embryonic root formation in Arabidopsis. A loss-of-function mutation in *Atarf5*/*mp* results in a rootless phenotype. AtARF5/MP directly activates several bHLH families of transcription factors which are a critical determinant of root development. Among these factors, *TARGET OF MONOPTEROS5 (TMO5)*, *TMO7,* and *LONESOME HIGHWAY (LHW)* is particularly important. *TOM5* and *LHW* have overlapping expression patterns and form heterodimers that activate the expression of *LOG4*, thereby establishing an auxin-mediated CK gradient. TOM7, on the other hand, is a small HLH protein that lacks a basic DNA binding region and has been suggested to function as an inhibitor. TOM5 and TOM7 do not show overlap in their expression domains. In the developing vascular tissue, a zone of high auxin and a surrounding zone of high CK govern vascular patterning. Thus AtARF5/MP and its bHLH targets (*TMO5*, *TMO7,* and *LHW*) mediate an intricate interplay of auxin and CK in a spatiotemporal manner leading to central vascular bundle development during root initiation (Ohashi-Ito and Fukuda, 2010; De Rybel et al., 2016). The role of MtARF5 in embryonic root development or factors that govern *M. truncatula* embryonic root development is unknown. This study has demonstrated that MtARF5 is controlling nodule vascular bundle development by recruiting *NVD1* (Fig. 8). The expression of *NVD1* is widespread in the root and increases significantly during nodule development, while, *NVD2* is more or less specifically associated with nodule development (Supplemental Fig. 1 and 5). Both *nvd* mutants exhibited normal root development. NVD1 and NVD2 belong to a legume-specific branch suggesting MtARF5 mediated recruitment of legume-specific modules during nodule vascular bundle patterning (Supplemental Fig. 2 and 4). Interestingly, the ortholog of *AtTMO3*, which encode an AP2-type transcription factor is among the DEGs in *nvd1* nodule (Supplemental table 1).

**Figure 8.**
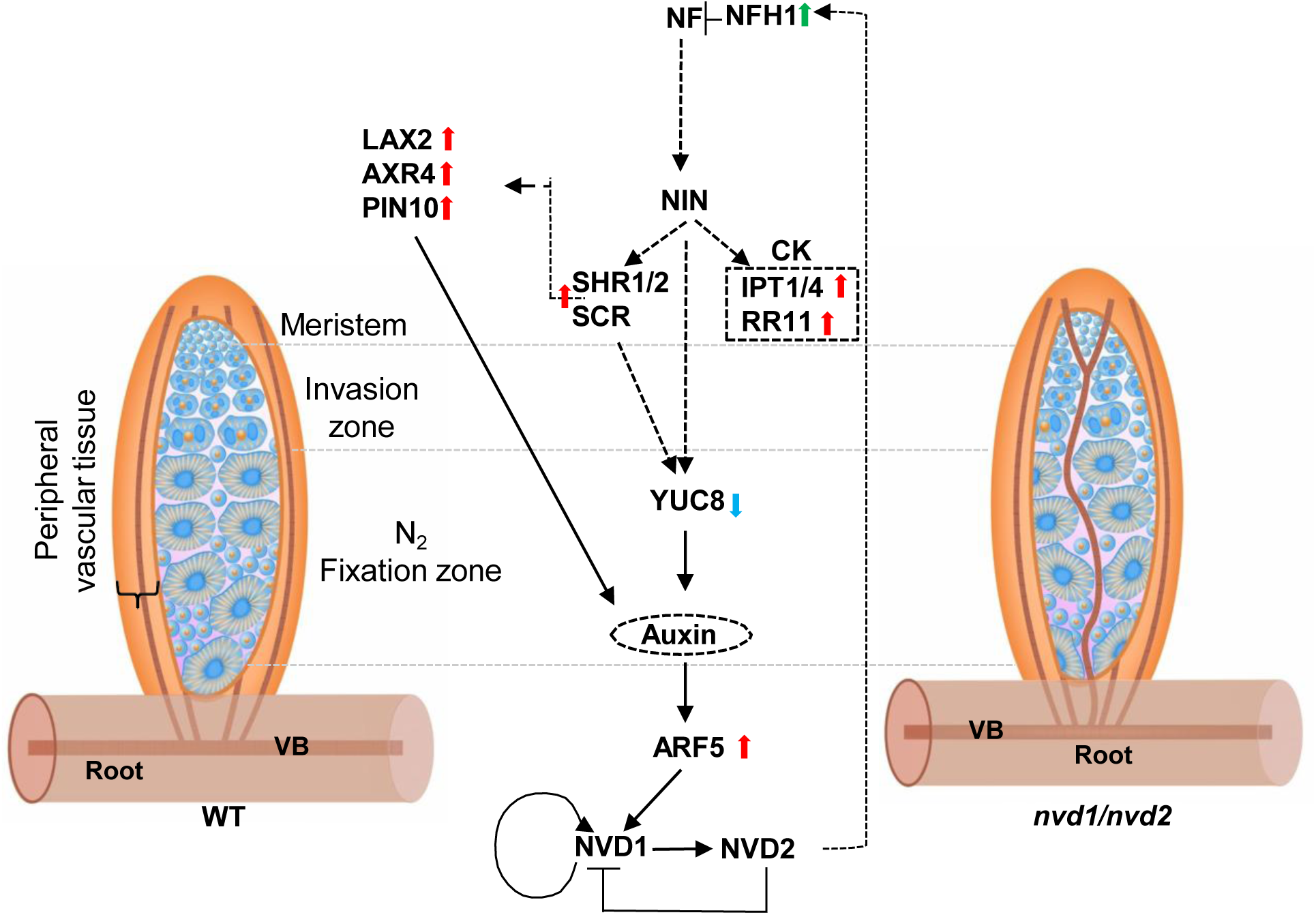
Schematic representation of a proposed signaling pathway that controls peripheral vascular bundle development in *M. truncatula* nodule. Auxin and MtARF5 activate *NVD1* expression. NVD1 activates *NVD2* and also its own expression in a feed-forward loop. NVD2 interacts with NVD1 and forms a non-functional heterodimer and thereby modulating NVD1 and NVD2 expression. *nvd1* nodules show elevated expression of *NFH1,* whereas both *nvd1* and *nvd2* nodules show elevated expression of *SHR/SCR*, *LAXs, AXR4,* and *PINs* and downregulation of *YUC8* thereby disrupting auxin distribution pattern. Cytokinin biosynthesis gene *IPT1/4* and type A response regulator *RR11* show elevated expression. Together auxin and CK disequilibrium resulted in distorted vascular bundle phenotype in *nvd1* and *nvd2* nodules. Solid lines indicate direct interaction. Red and blue arrows indicate elevated expression and downregulation in both *nvd1* and *nvd2* nodules. The green arrow represents elevated expression only in *nvd1* nodules.

NF signaling leads to the development of nodules with peripheral vasculature, but how the latter is achieved when the default for roots is a central vasculature remains unclear. Our results add some clarity to this question. NF signaling activates CK transport from the epidermis to the cortex/pericycle (Jarzyniak et al., 2021). Subsequently, CK activates *NIN* expression in root pericycle cells (Liu et al., 2019). The NIN signaling pathway activates biosynthesis of auxin ∼10-12 hpi in *M. truncatula* roots and promotes cell division ∼24 hpi (Schiessl et al., 2019). Constitutive activation of the NF-signaling pathway leads to the formation of spontaneous nodules with an ontology, including peripheral vasculature similar to the WT nodules (Gleason et al., 2006; Tirichine et al., 2006b; Tirichine et al., 2006a; Tirichine et al., 2007; Ried et al., 2014; Saha et al., 2014). In contrast, when the NF signaling pathway is manipulated downstream of the early CK signaling the resultant nodule-like structures have a central vasculature. Likewise, exogenous application of auxin efflux transport inhibitors to roots or perturbation of the CK signaling by overexpressing *MtKNOK3* promotes the formation of nodule like structure with a central VB (Soyano et al., 2013; Azarakhsh et al., 2015; Kohlen et al., 2018; Soyano et al., 2019; Dong et al., 2021). Thus, as opposed to auxin govern CK biosynthesis during lateral root development, CK-mediated recruitment of auxin signaling promotes peripheral vascular bundle development. *MtNFH1*, which encodes a hydrolase that degrades NF (Cai et al., 2018) is upregulated in *nvd1* nodules and presumably interferes with NF-signaling. Our transcriptome and microscopic data indicate that the auxin/CK balance is disturbed in the *nvd* nodules (Fig 6 and 7). Upregulation of *SHR2*, *LAX2, AXR4, ARF5, PIN10,* IPT1/4, RR11, and orthologs of Arabidopsis vasculature development-related genes *MtLITTLE ZIPPER*, *MtJACKDAW* in both *nvd* nodules indicated that the nodule developmental program becomes more like the root development. Effective repression of root identity by NOOT1/2 is important for normal nodule development, while increases expression of *NOOT1* and *NOOT2* in both *nvd* nodules indicate an overly active root developmental program. Thus, the crosstalk between NF and hormonal signaling appears to be disrupted in *nvd* nodules.

Downstream of the NF-signaling, CK activates *NIN* and the NIN signaling pathway activates auxin biosynthesis in the root tissue. During *M. truncatula* nodule development, auxin biosynthesis gene *YUCs* and auxin influx carrier *LAX2* are activated around 12-16 hpi (Schiessl et al., 2019). *MtARF5* and *NVD1* induction coincided with the auxin maxima (Supplemental Fig. 1B). Induction of *NVD1* ∼ 14-16 hpi indicated that priming of development of multiple peripheral vascular strands before the onset of cell division in the root pericycle. In a comparative study between *M. truncatula* lateral root and nodule development, it was shown that lateral root development is much faster, and the central vascular strand appears earlier compared to nodules (Schiessl et al., 2019). Restriction of rhizobial infection to the root epidermis due to a mutation of either the plant genes or the bacterial gene *exoY* leads to the formation of a central vascular bundle in *M. truncatula* nodules (Guan et al., 2013). Auxin signaling is activated in the root cortex after inoculation with rhizobial *exoY* mutants. In contrast, auxin signaling is restricted to the pericyclic cell layers when plants are inoculated with wild-type bacteria (Guan et al., 2013). Thus, a change in spatiotemporal auxin signaling seems to be a prerequisite for peripheral vascular bundle development. We found perturbation of several auxin and CK signaling pathway genes and altered auxin/CK distribution patterns in *nvd* nodules (Fig. 6 and 7), implicating NVD1/NVD2 in the control of auxin and CK balance in a feedback loop (Fig. 8). NVD1 work in a feedforward loop to control its expression while NVD1 is regulated at the protein level by NVD2 (Fig.3C-F and Fig.7C). NVD1 directly activate *NVD2*. According to the nodule developmental time course, NVD1 started expressing ∼ 14 hpi whereas *NVD2* gets activated at ∼36 hpi. Of note, the *pNVD2* contains multiple Auxin-Responsive Elements (*Aux-REs*) and can be activated in a transactivation assay by MtARF5. However, NVD2 is not induced in nodules lacking *NVD1*, indicating that NVD1 is essential for *NVD2* activation in *M. truncatula*. This is also consistent with the timing of nodule development, where the auxin maxima coincide with *NVD1* expression, but *NVD2* expresses much later in the time scale (Fig. 7 and Supplemental Fig. 1 and 5). As NVD2 started building up in a developing nodule primordium it will control both *NVD1* transcription as well as NVD1 activity (Fig. 3). The complexity involved in determining the appropriate dosage of NVD1 and NVD2 at a spatiotemporal context is evident from the multiple parallel findings highlighted above (Fig. 8). Additionally, despite the broader expression, the significance of specific localization of NVD2 in the NVE is not clear (Fig. 5), but this data suggest NVD2 might also be regulated at the translational level. In *nvd1* nodules, *MtNFH1* gets upregulated. MtNFH1 degrades NF and thereby modulates the NF-signaling pathway. Downregulation of *NIN*, *NSP2,* and *EFD1* in *nvd2* nodules could be due to the perturbation in the NF-signaling pathway (Fig. 8). The NIN signaling pathway activates *YUCs* during nodule development (Schiessl et al., 2019; Shrestha et al., 2020). Therefore, the downregulation of *MtYUC8* in *nvd* nodules could also be explained by NFH1-mediated degradation of NF (Fig. 8). Despite the downregulation of *YUC8* we found upregulation of *LAX2* and four *ARFs* in *nvd* nodules, along with altered auxin distribution. In Arabidopsis, auxin influx carrier *AtLAX3* expression is regulated by AtSHR (Sozzani et al., 2010; Della Rovere et al., 2015). Further, AtSHR/SCR and AtAUX1 control the auxin balance to promote adventitious rooting or xylogenesis in Arabidopsis (Della Rovere et al., 2015). In *nvd* nodules, we have noticed simultaneous activation of *MtLAX2, MtAXR4,* and *SHR*/SCR. Thus, an alternative pathway(s) is operational in absence of *NVD1* or *NVD2,* which controls auxin distribution inside the nodule primordium leading to the spatiotemporal changes in the auxin distribution pattern. In this connection, EFD1 activates *MtRR4* during nodule development. In the *nvd2* nodule, *EFD1* is downregulated while *MtRR4* is upregulated, highlighting alternative pathways that take over CK signaling in *nvd2* nodules. Noteworthy, over-expression of *MtRR9* resulted in the formation of arrested primordia, containing key developmental programs necessary for the formation of both nodules and lateral roots (Op den Camp et al., 2011). In both *nvd1* and *nvd2* nodules another NF-induced type A response regulator MtRR11 is overexpressed (Fig. 6E). Thus, fine-tuning of the CK and auxin signaling appears to have become imbalanced in *nvd* nodules. This is also evident by the output of auxin and CK signaling at the nodule apex (Fig. 7 G-K). We propose, that this is the primary factor contributing to the abnormal development of peripheral vascular bundles (Fig. 8).

A single point of predisposition led to the recruitment of genes needed for root nodule developmental traits in the nitrogen-fixing clade (NFC) (Werner et al., 2014; Griesmann et al., 2018). However, during evolution, the vascular bundle shifted to the periphery of the nodule in legumes. Previously, it has been proposed that legumes adopted peripheral vascular bundles by establishing fine coordination between infection and organogenesis. In contrast, nodule-forming non-legumes separate epidermal infection and nodule organogenesis spacio-temporally (Guan et al., 2013). Our study highlighted that NVD1 and NVD2 along with a fine balance of auxin/CK is necessary for the establishment and maintaining the orderly arrangement of peripheral nodule vasculature. We discovered the presence of a complex regulatory circuit with multiple layers of redundant regulations behind the peripheral vascular bundle development. The Macroscopic phenotype is only present in ∼30 % of nodules in *nvd1* plants, and in *nvd2* the phenotype is even weaker (Compare Fig. 1 and Fig. 4). We observed the overexpression of several bHLH-TFs in *nvd* nodules (Supplemental Fig. 6E), which might be the reason behind the low penetrance of the phenotype due to the redundant action of bHLH-TFs. Together, our findings indicate that robust and redundant machinery controls peripheral vascular bundle development in legumes.

In summary, the unique phenotype of *nvd1* and *nvd2* enables us to understand how peripheral vascular bundle development is tightly controlled in legumes. Compromised nitrogen fixation in *nvd* nodules highlights the physiological advantage of having a vascular bundle in the nodule periphery. The signals triggering the peripheral vasculature development or if that is a single determinant remain elusive. The connections among *NVD1/2*, *NOOT1/2*, NF signaling, and lateral root developmental programs also need to be explored in the future. *NVD1*, *NVD2,* and the pathway that recruits *NVD1/2* during nodule development are crucial determinants of efficient nitrogen fixation by legume nodules. Single nucleotide polymorphisms associated with these genes can be used for a breeding program in the future to obtain high nitrogen-fixing nodules.

## Methods

### Plant growth and bacterial strain

*M. truncatula* ecotype R108, three *nvd1* (MtrunA17_Chr3g0132471) mutant lines *nvd1-1*(NF4649), *nvd1-2* (NF13338) *nvd1-3* (*NF5171*) and two *nvd2* (MtrunA17_Chr5g0437991) mutant*s nvd2-1*(NF19150)*, nvd2-2* (NF7653) were used. Seeds were scarified for 8 minutes in sulfuric acid followed by washing five times with autoclaved water, followed by surface sterilization using 30% (v/v) commercial bleach containing 4% (v/v) tween 20 for 8-10 minutes and washed thoroughly with autoclaved water until the smell of bleach faded off. Seeds were kept at 4°C in dark for 1 day. Seeds were plated in sterile petri-plate having damp, sterile filter paper and kept inverted at 20-23°C in dark for 1 day for radicle growth and then in the day light cycle for 2 days. Seedlings were transferred to soil (Soilrite, Keltech Energies Ltd.). Plants were cultivated at 21°-23°C under photoperiod of 16-h light/8-h dark with 250 µE.m^-2^s^-1^ light irradiance and 40% relative humidity.

Plants were inoculated with *S. meliloti* strain 1021after 7-10 DAS. *S. meliloti* were grown for 20-24 hrs in Tryptone Yeast (TY) media at 28°C till OD_600_ reaches ∼ 1.0 and harvested. For inoculation to plants, bacterial pellets were resuspended to B&D medium (Broughton and Dilworth, 1971) to OD_600_ ranges from 0.03 - 0.05 and 50 ml of inoculum was added to each pot. For hairy root transformation *Agrobacterium rhizogenes* ARqua1 and for *N. benthamiana* transient expression *A. tumefaciens* strain GV3101 were used.

### Constructs

The primers used for PCR amplification are shown in supplemental table 3. For transactivation assay *NVD1*, *ARF5* were amplified from *M. truncatula* (R108) cDNA and cloned in *pENTR-D-TOPO* vector (Invitrogen) and recombined in *pK7FWG2* (Karimi et al., 2002) using LR clonase (Invitrogen). *pNVD1^2kb^* and *pNVD2^1kb^* were amplified from the *M. truncatula* (R108) genomic DNA and cloned into *pENTR-D-TOPO* (Invitrogen) and recombined into *pKGWFS7* (Karimi et al., 2002) upstream to *GUS* using LR clonase (Invitrogen). Promoter fragment *pNVD2^1kb^* and *NVD2^185bp^*recombined in *pKGWFS7* were used for transactivation assay. Further *pNVD2^185bp^*was cloned in *pKGWFS7* for promoter reporter assay. For Y1H assay, *NVD1* is recombined in *pGADT7* through gateway cloning strategy and *pNVD2^185bp^*in *pAbAi* using traditional cloning method. For split luciferase complementation assay, *NVD1* and *NVD2* were cloned in *pCAMBIA1300-C_Luc* and *N_Luc* vectors (Zhou et al., 2018) respectively through restriction-based cloning. *AtBIN2* cloned in *pCAMBIA1300 N_Luc* was used as negative control. For ectopic expression, *NVD2* was cloned into *pKGW-RR-ENOD12-cGFP* (*Sinharoy et al., 2016*) using LR clonase (Invitrogen). For subcellular localization studies *pNVD2^1kb^* was fused with *NVD2* using overlapping PCR and recombined in *pKGW-RR-cGFP* via gateway cloning.

### Hairy Root Transformation of *Medicago truncatula* R108

Composite plants was generated as described in (Sinharoy et al., 2015). Briefly*, A. rhizogenes* (ARqua1) strain containing a desirable construct was grown overnight in TY media. A bacterial lawn was prepared on a media plate with antibiotic selection. Seedlings with a small excision at the tip of radicle were scraped to bacterial plate and placed to Fahraeus media plates. The composite plants were transferred to pots after 7-10 days and inoculated with *S. meliloti*.

### Acetylene Reduction Assay

Acetylene reduction assay (ARA) were performed at 10, 15 and 21 dpi as described previously (Oke and Long, 1999) for *nvd1* and at 45 dpi for *nvd2*. For ARA the entire root was separate from the shoot part, placed in a tube and sealed with suba-seal septa (sigma-aldrich). 1 ml of pure acetylene was injected through suba-seal septa and incubated in dark at room temperature for 16 hours. Ethylene production was measured by gas chromatography (Shimadzu GC-2010 equipped with HP-PLOT ‘S’ Al_2_O_3_ 50m,0.53mm column (Agilent Technologies) as described in (Mandal and Sinharoy, 2019).

### RNA isolation and qRT-PCR analysis

Total RNA was isolated using RNA purification kit (Nucleospin RNA Plant, Macherey-Nagel, Düren, Germany) and treated with DNase (Turbo DNA free kit, Thermo Fisher). One microgram purified RNA used for cDNA synthesis using Verso cDNA synthesis kit (Thermo Fisher). Quantitative reverse transcriptase PCR (qRT-PCR) was performed using PowerUp SYBR Green master mix (Thermo Fisher) on a CFX96 Touch Real-Time PCR Detection System (Bio-Rad). The transcript level of indicated genes were normalized using ubiquitin conjugating enzyme 9 (*UBC9*) (MtrunA17Chr7g0275361). The primers used are listed in supplemental table 3.

### Yeast One Hybrid Assay

Y1H Gold strain was transformed with *pAbAi* containing 185 bp region of *pNVD2* using EZ-Yeast transformation kit (MP Biomedical). Positive colonies were sequentially transformed with *pGADT7* containing *NVD1*. Positive yeast colonies were confirmed by growing on synthetic dropout media with - Leu and -Ura (LU) plates. Confirmation of positive interaction was done by growing the yeast colonies in dilutions of 1/5, 1/10 and 1/25 on -LU plates with Aureobasidin (150 ng/ml concentration) as described in Matchmaker Gold Yeast one-hybrid user manual.

### Transactivation and split luciferase complementation assay

*A. tumefaciens* strain GV3101 carrying *pNVD2_1kb_::GUS*, *pNVD2_185bp_::GUS, p35S::NVD1:GFP* or empty vector (*pK7FWG2*) were infiltrated into *N. benthamiana* leaves along with *P19* containing plasmid (Angel et al., 2011) to inhibits gene silencing. Plants were kept in dark for 12 hours and transferred in light for 24-36 hours. For histochemical GUS visualization, infiltrated leaves were incubated for 12 h at 37°C in the GUS-staining solution containing 0.1 M sodium-phosphate buffer (pH 7), 10 mM EDTA, 0.1% [v/v] triton X-100, 1 mM potassium Ferricyanide, and 1 mg/ml 5-bromo-4-chloro-3-indoxyl-β-D-glucuronide cyclohexyl ammonium salt (X-Gluc) (Gold Bio). Leaves were photographed after chlorophyll removal using 70 % ethanol. For quantification qRT-PCR was done using *GUS* and *NPTII* (as an internal control) with primer sequences listed in Supplemental table 3. For split luciferase complementation assay, infiltration was done using *A. tumefaciens* GV3101 in *N. benthamiana* leaves. LUC images were taken 2-3 days after infiltration by applying 1 mM Beetle luciferin (Promega) on adaxial leaf surface.

### Transcriptome analysis

15 dpi root nodules were chopped and collected for RNA sequencing (RNA Seq) for R108, *nvd1-2, nvd1-3, nvd2-1* and *nvd2-2*. Total RNA was isolated using QIAzol Lysis Reagent by the manufacturer’s protocol. Strand-specific mRNA libraries were prepared using the Breath Adapter Directional sequencing (BrAD-seq) method (Townsley et al., 2015). RNA-seq was performed using Illumina HiSeq platform. The reads were quality filtered using NGSQC toolkit that involved removing low-quality reads (Phred score < 30) and adapter sequences. Two independent *Tnt1* insertional lines of *nvd1* and *nvd2* were used for RNAseq. As *Tnt1* mutants may have 25-100 background insertions (Tadege et al., 2008; Sun et al., 2019), hence this strategy was adopted to reduce any interference due to background mutation. The use of two different insertional lines may have reduced the number of DEGs in our analysis, but we obtained a robust dataset. Two biological and two technical replicates were used for analysis. Approximately 4GB of clean reads were obtained for each sample. *S. meliloti* 1021 sequences were removed using the Sortmerna tool. The remaining reads were indexed and mapped with the *M. truncatula* A17 mRNA v.5.0 (https://medicago.toulouse.inra.fr/MtrunA17r5.0-ANR) using the default parameter of Bowtie 2. Approximately 65% data was mapped with the reference mRNA. Out of the total number of 44,618 genes available in reference mRNA, reads were mapped to a minimum of 22,804 genes from different libraries. The raw read count was normalized and differential expression analysis was performed using DESeq2 package in R programming environment. The criteria of Log2 fold change ≥1 (upregulated genes) and ≤ -1 (downregulated genes) and adjusted P-value cut-off < 0.05 were used for determining significant differentially expressed genes.

### Phylogenetic analysis

Tree available in the Arachis hypogaea Nodule Developmental Gene Expression (AhNGE - http://14.139.61.8/AhNGE/index.php) were downloaded for NVD1 and NVD2. These trees were built during the orthofinder analysis and the visualization of the trees was done using Interactive Tree of Life (iTOLv5)(Letunic and Bork, 2007).

### Phenotypic analysis, GUS staining and microscopy

Fresh weights of R108, *nvd1* and *nvd2* were taken using a weighing machine (Sartorious). For nodule length measurement and nodule counting plants were scanned. ImageJ software was used for nodule length measurement and nodule number counting manually. Images of whole nodules and sections were captured using Nikon stereoscopic zoom microscope SMZ25. Transformed roots with the promoter-GUS construct were stained in a solution containing 0.1 M sodium-phosphate buffer (pH 7), 10 mM EDTA, 0.1% [v/v] Triton X-100, 1 mM potassium ferricyanide, and 1 mg/ml 5-Bromo-4-chloro-3-indoxyl-β-D-glucuronide cyclohexyl ammonium salt (X-Gluc) (Gold Bio). Samples were vacuum infiltrated and incubated at 37°C. Detached nodules were fixed for 1 hour in 3% paraformaldehyde (w/v) and 2.5% glutaraldehyde (v/v) in 0.2 M sodium cacodylate buffer (pH 7.2). The samples were washed with 0.05 M sodium cacodylate buffer and embedded in 4% agarose and sliced using VT1000S vibratome (Leica). For localization and auxin/cytokinin distribution pattern, nodules were hand sectioned and confocal microscopy was performed using Leica TCS-SP8 microscope (Leica microsystems, Wetzlar, Germany). Images were captured at 488nm excitation (ex) and 500-530 nm emission (em) for GFP; 561 nm ex and 570-600 nm em for TdTomato. The images were processed using Photoshop CS6X64.

### Auxin treatment

A 10 mM IAA (Sigma) stock solution was prepared in 50%-water, and 50%-ethanol. 10 µM and 20 µM working stocks of IAA were prepared in water. Plants were grown on Farhaeus media plates containing for 15 days. 5 ml of 10 µM solution was applied on the roots of 15 days old R108 seedlings and for *nvd1* seedlings, 20 µM solution was treated for 12 hours.

### Live/Dead staining of rhizobia

15 dpi nodules were hand sectioned and kept for 10 minutes in staining solution (1:3 ratio of 1:500 dilution of propidium Iodide (Sigma) and 1: 500 dilutions of SYTO 13 (Life Technologies) in water. The section was removed from the staining solution and mounted in deionized water for visualization using Leica TCS-SP8 microscope (Leica microsystems, Wetzlar, Germany).

## Author Contributions

SS and AR designed the experiments. DS, AG, and SS performed experiments, analyzed data, and prepared the figures. SS, DS, and MU wrote the manuscript.

## Acknowledgments

We thank S. Subramanian, South Dakota State University, USA, for 2-COL and AP Singh for the BIN2 cloned in pCAMBIA1300 N_Luc. We acknowledge NIPGR for confocal facilities; CIF-NIPGR and DBT (Department of Biotechnology)-eLibrary Consortium (DeLCON), India, for providing access to e-resources. Work was supported by research grants from NIPGR. DS was supported by CSIR-SPM (07/803(0329)/2020-EMR-I).

## Supplement Figures

**Supplement Fig 1.**
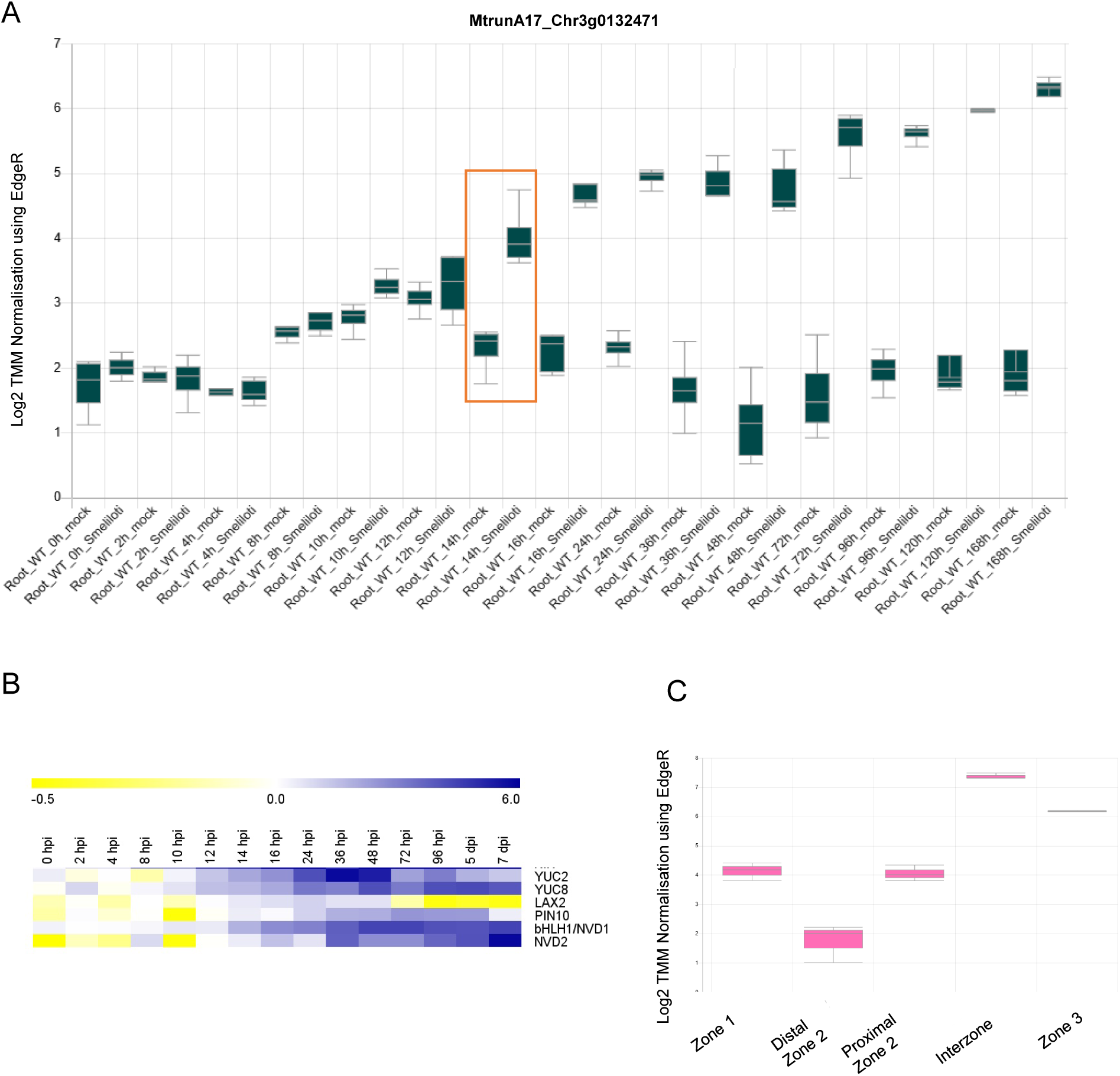
Expression profile of *NVD1* gene in the inoculated root, different nodule zone, and its time course induction in correlation with the other genes. (A) *NVD1* gene expression from 0 to 168 hours post inoculation (hpi) was analyzed using MtExpress. *NVD1* has induced 14 hours post inoculation (hpi) (as indicated by the red box) and continued up to 168 hpi. (B) Heatmap showing gene expression as published by (Schiessl et al., 2019) for *NVD1*, *NVD2,* and auxin-related genes that are differentially expressed in *nvd1* and *nvd2* nodules. (C) *NVD1* expression in different nodule zones highlighted the highest abundance in NVD1 expression in the Interzone and nitrogen fixation zone, according to the MtExpress tool (Carrere et al., 2021).

**Supplement Fig. 2.**
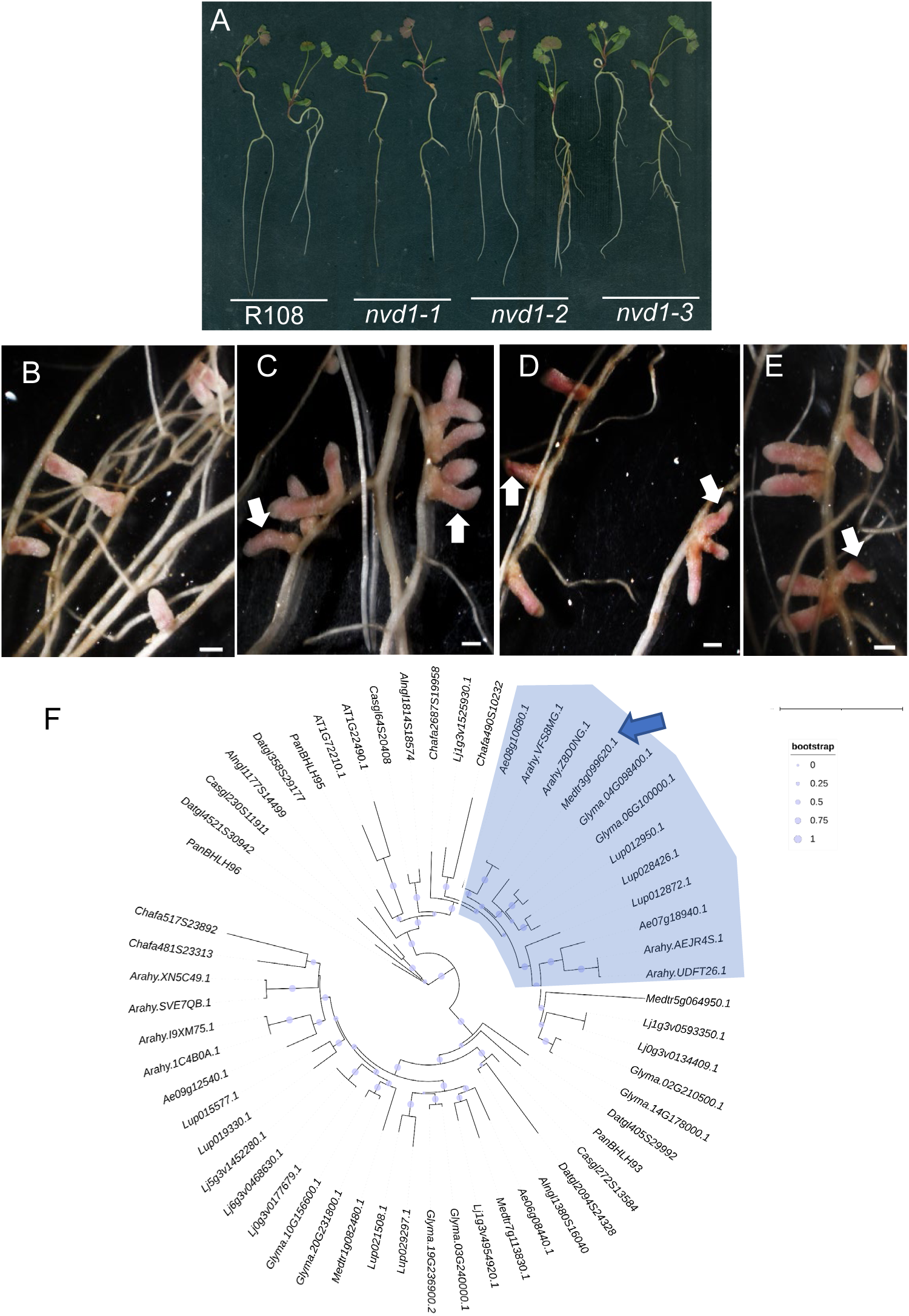
Phenotype of *nvd1* nodules and phylogenetic analysis of NVD1. (A-D) Images of R108 and *nvd1* nodules formed at 30 days post-inoculation (dpi). R108 (A), *nvd1-1* (B), *nvd1-2* (C) and *nvd1-3* (D). All plants form pink-colored nodules. (E) Phylogenetic analysis of NVD1 and its orthologs from legumes and non-legumes. The highlighted area in blue shows NVD1 and its orthologs from legumes clustered together. An arrow represents the MtNVD1. Scale bar = 1 mm (A-D).

**Supplement Fig 3.**
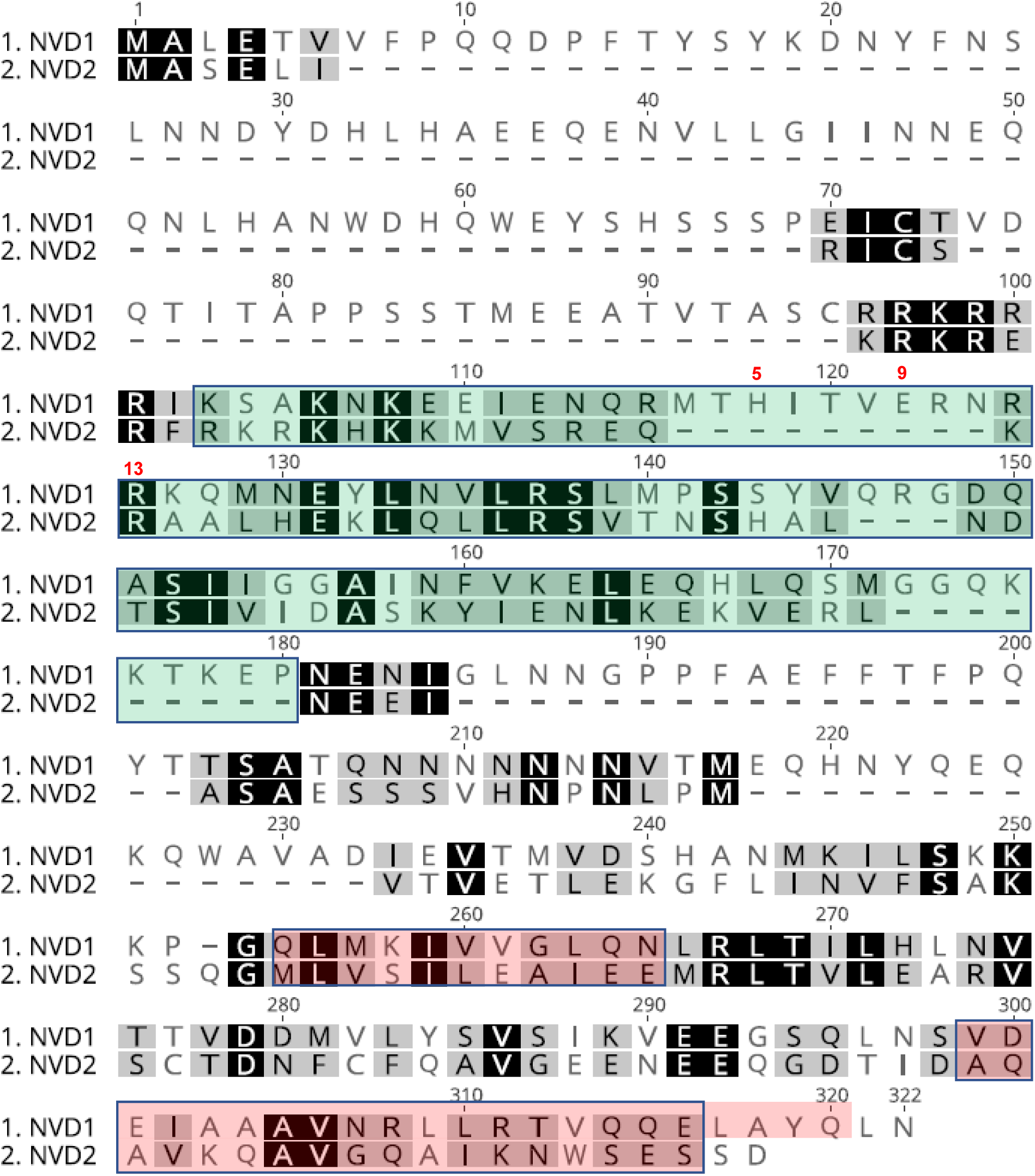
Alignment of NVD1 and NVD2 sequences. Sequence alignment of NVD1 and NVD2 showing conservation of the Helix Loop Helix (Protein-Protein Interacting) domain in NVD1 and NVD2, while NVD2 lacks the basic (DNA Binding) domain. The green and pink coloured box represents the DNA Binding and Helix Loop Helix domains respectively.

**Supplement Fig 4.**
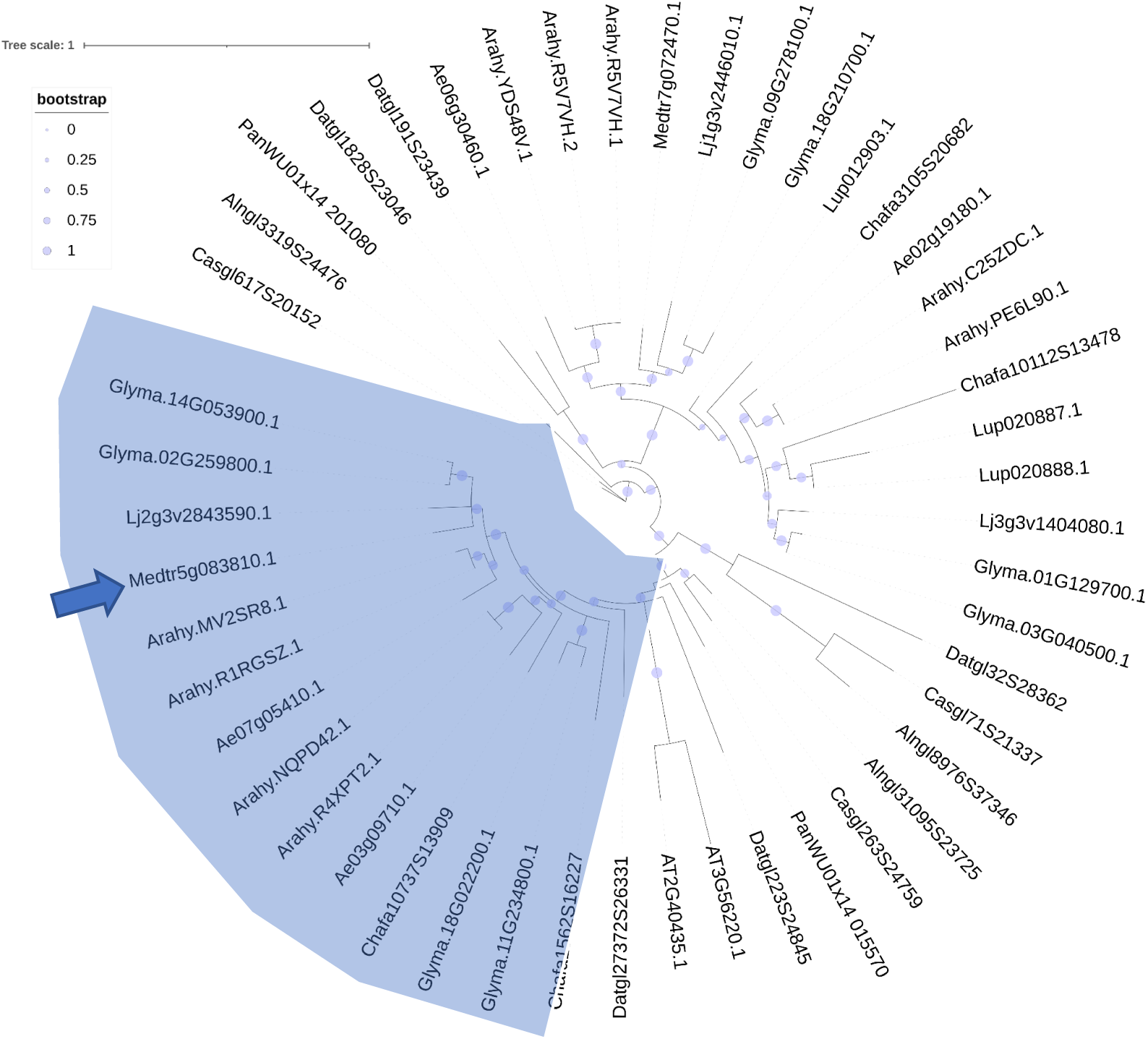
Phylogenetic analysis of NVD2. A phylogenetic tree of NVD2 using orthologous proteins from legumes and non-legumes showing NVD2 from legumes getting clustered together (highlighted in blue). An arrow represents the position of MtNVD2.

**Supplement Fig 5.**
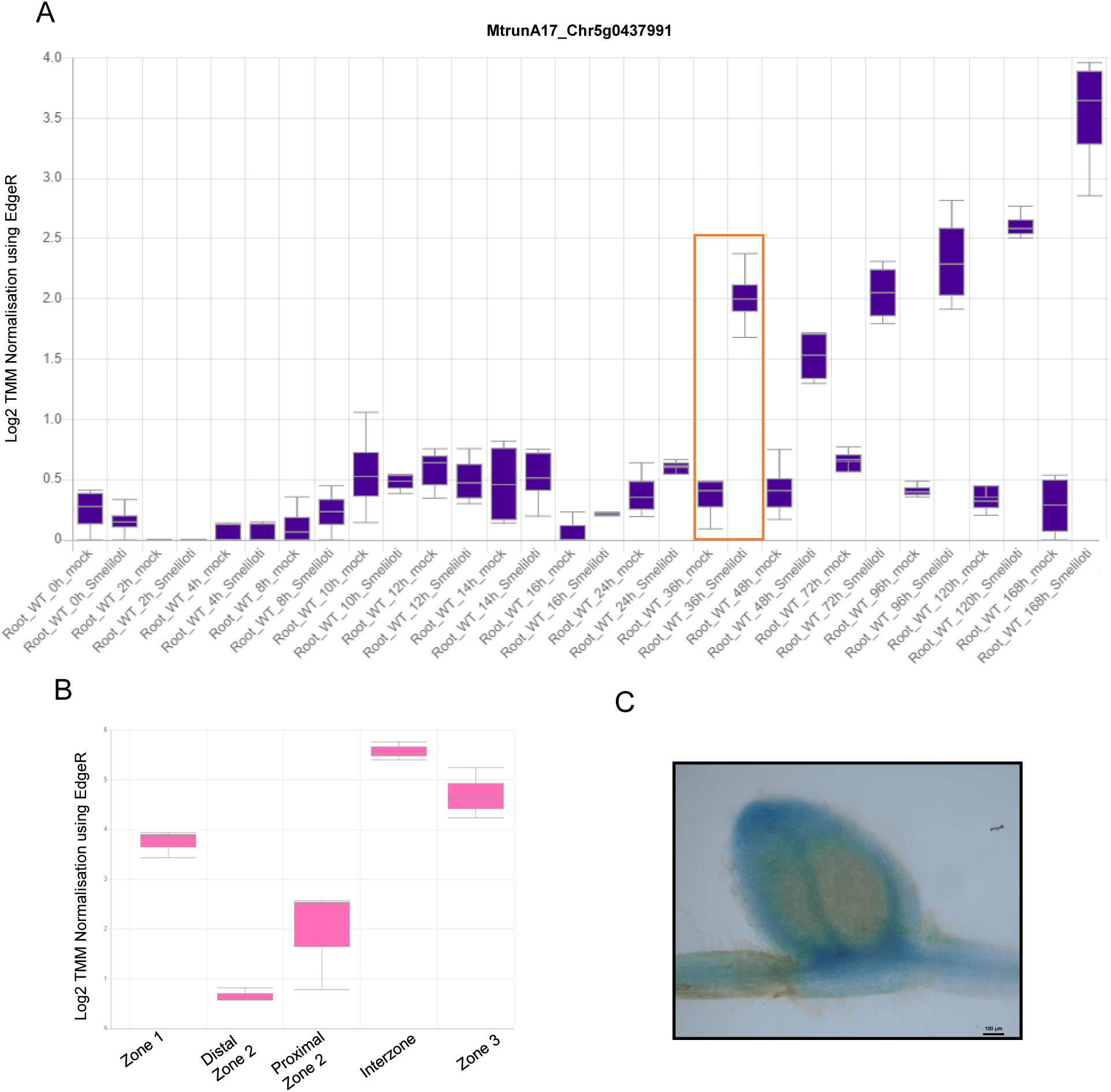
Expression profile of the *NVD2* in inoculated root and different nodule zones. A) Expression profile of the *NVD2* in inoculated root from 0 hpi to 168 hpi in accordance with the MtExpress. *NVD2* shows significant induction at 36 hpi (represented by the red box). B) *NVD2* expression in different nodule zones with the highest abundance in the invasion and nitrogen fixation zone according to the MtExpress tool (Carrere et al., 2021).

**Supplement Fig 6.**
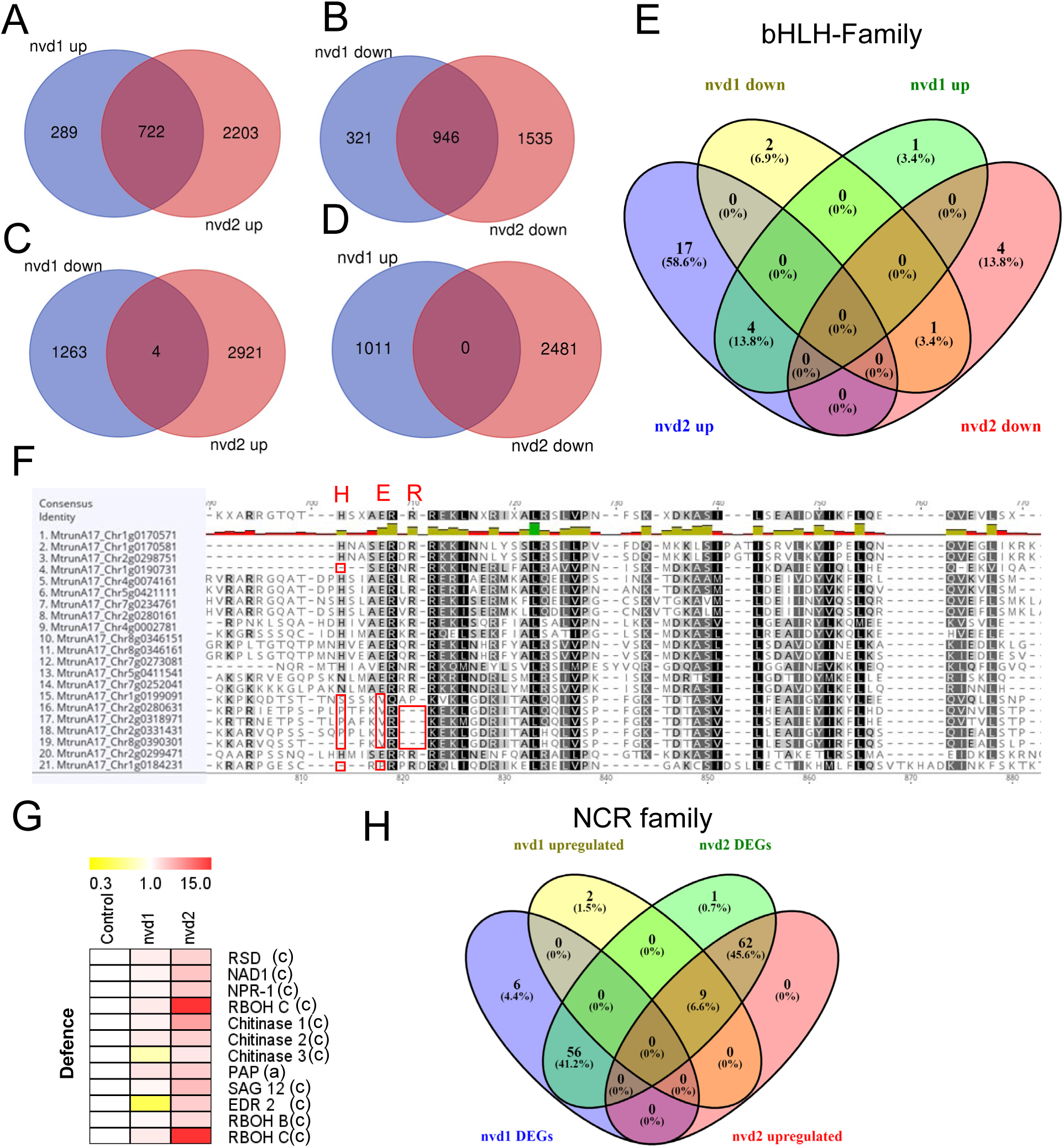
Comparison of the DEGs of *nvd1* and *nvd2* nodules. (A-D) Venn diagram showing DEGs in *nvd1* and *nvd2* nodules (A), upregulated (B), downregulated (C-D), differential expression in opposite direction in *nvd1* and *nvd2* nodules. (E) Venn diagram showing the comparison of total differentially expressed bHLH transcription factor (TF) versus upregulated TF in *nvd1* and *nvd2* nodules. 21 *bHLH TF* are upregulation in *nvd2* nodules. (F) Alignment of upregulated bHLH-TF sequences in *nvd2* nodules; 7 out of 21 bHLH-TFs lack basic DNA binding regions and are hence considered as HLH proteins. Changes in crucial amino acid residues in DNA-binding domain or the absence of amino acids in DNA-binding domains are highlighted with red boxes. (G) Heatmap showing the upregulation of *RSD*, *NAD1*, senescence, and pathogenesis-related genes in *nvd2* nodules. (H) Venn diagram represents differentially expressed *NCRs* in *nvd1* and *nvd2* nodules. Among 120 *NCRs*, 68 *NCRs* get upregulated in *nvd2* nodules.

**Supplement Fig 7.**
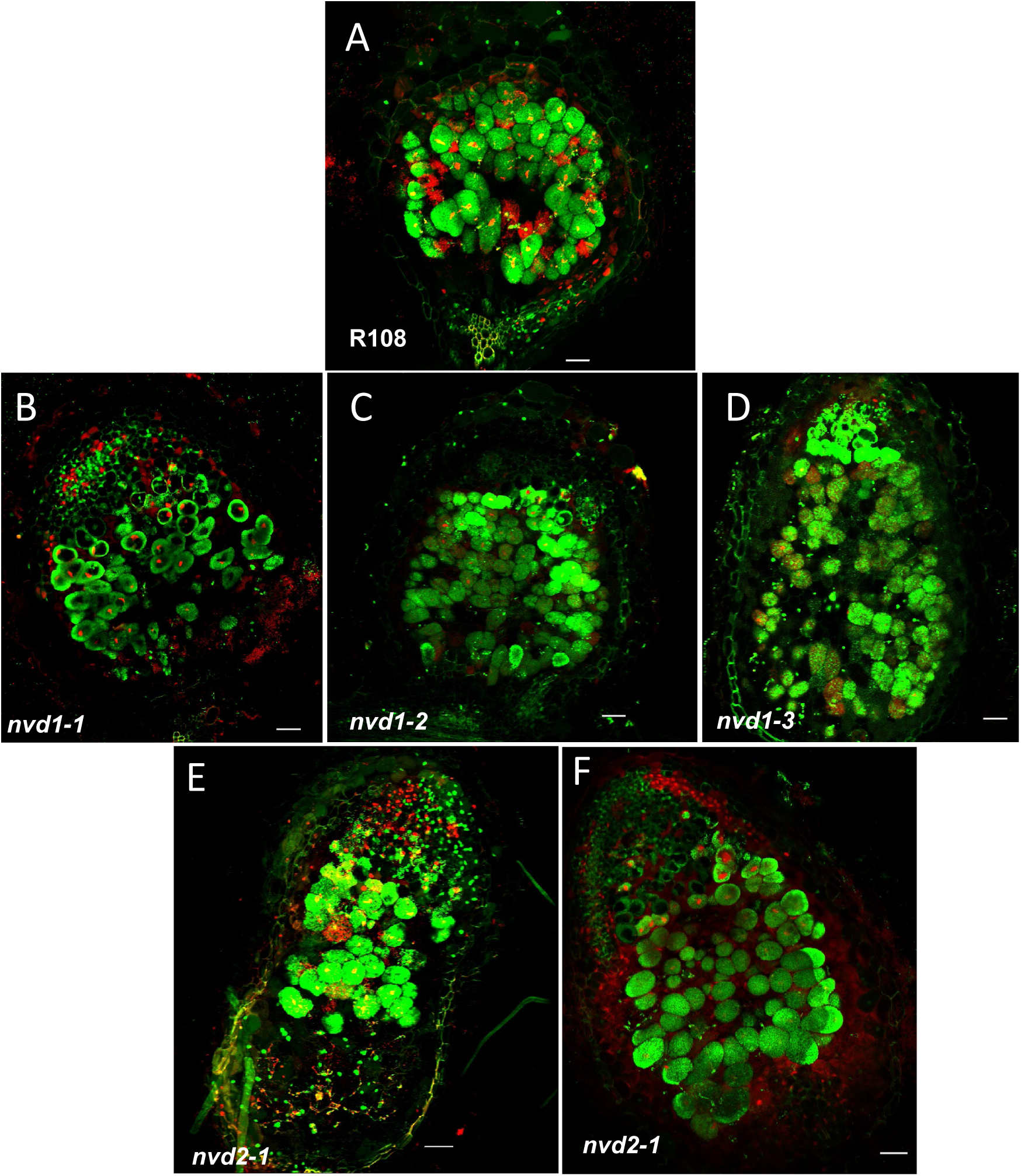
Live/dead staining of R108, *nvd1,* and *nvd2* nodules. (A-F) Maximum intensity projection of merged confocal images of R108 (A), *nvd1* (B-D), and *nvd2* (E, F) nodules at 15 dpi. Nodules were stained with SYTO 13 (Green) and Propidium Iodide [PI] (Red). Live bacteria are stained with SYTO13 and dead bacteria are with PI. Scale bar = 50 µM.

## Supplemental Tables

Supplemental Table 1: Differentially expressed genes in the *nvd1* nodules at 15 dpi.

Supplemental Table 2: Differentially expressed gene involved in auxin and cytokinin biosynthesis and signaling pathway.

Supplemental Table 3: Primers used for this study.

## Notes

### Competing Interest Statement

The authors have declared no competing interest.

